# Evaluating genomic data for management of local adaptation in a changing climate: A lodgepole pine case study

**DOI:** 10.1101/568725

**Authors:** Colin R. Mahony, Ian R. MacLachlan, Brandon M. Lind, Jeremy B. Yoder, Tongli Wang, Sally N. Aitken

## Abstract

The need for tools to cost-effectively identify adaptive variation within ecologically and economically important plant species is mounting as the detrimental effects of climate change become increasingly apparent. For crop and wild populations alike, mismatches between adaptive variation and climatic optima will reduce health, growth, survival, reproduction, and continued establishment. The ease with which land managers can quantify the relative importance of different climate factors or the spatial scale of local adaptation to climate will have direct implications for the potential of mitigating or resolving such risks. Using seed collected from 281 provenances of lodgepole pine (*Pinus contorta*) from across western Canada, we compare genomic data to phenotypic and climatic data to assess their effectiveness in characterizing the climatic drivers and spatial scale of local adaptation in this species. We find that genomic and climate data are nearly equivalent for describing local adaptation in seedling traits. We also find strong agreement between the climate variables associated with genomic variation and with 20-year heights from a long-term provenance trial, suggesting that genomic data may be a viable option for identifying climatic drivers of local adaptation where phenotypic data are unavailable. Genetic clines associated with cold injury occur at broad spatial scales, suggesting that standing variation of adaptive alleles for this and similar species does not require management at scales finer than are indicated by phenotypic data. This study demonstrates that genomic data are most useful when paired with phenotypic data, but can also fill some of the traditional roles of phenotypic data in management of species for which phenotypic trials are not feasible.

## 1 Introduction

The impact of climate change is undeniable and particularly evident in forests of western North America. Evidence of tree injury and mortality from droughts, floods, wildfires, disease, and insect outbreaks is mounting rapidly (van Mantgem et al. 2009; Allen et al. 2010; Anderegg et al. 2015; McDowell & Allen 2015; Reyer et al. 2015; Buotte et al. 2018). There is also mounting evidence that changes in climate are disrupting local adaptation in plants (Mcgraw et al. 2015; Wilczek et al. 2019), with impacts to productivity of commercial tree species (Rehfeldt et al. 1999; Leites et al. 2012) and conservation of vulnerable species (Parmesan 2006). In response, forest managers are seeking guidance on which source populations to use for planting, as the long-practiced ‘local is best’ strategy no longer matches trees with the climates to which they are adapted (Aitken and Bemmels 2016). There is also a need to characterize the spatial scale and genetic structure of local adaptation to understand the capacity of populations to adapt to climate change without human intervention (McKenney et al. 2007; Kawecki 2008; Aitken et al. 2008; Kremer et al. 2012). For centuries, local adaptation has been quantified and managed using phenotypic data from long-term provenance trials and short-term common gardens (Langlet 1971; Leimu and Fischer 2008; Hereford 2009). In the past two decades, detailed climate data has been used to extend phenotypic inferences of local adaptation across managed landscapes (Sork et al. 2013; Wadgymar et al. 2017) and to project mismatches between adaptive variation and future climates (e.g., Exposito-Alonso et al. 2018). Genomic data is now emerging as a third source of insight into local adaptation for non-model species. While the genomic basis of local adaptation has been extensively studied (Li et al. 2017; Sork 2018), applications of genomic data to mitigate effects of climate change are in their infancy (Shafer et al. 2015). These applications can be advanced by understanding the ways in which genomic data complements and overlaps with phenotypic and climatic data in characterizing local adaptation.

For most tree populations, the capacity to track suitable climates via migration and establishment will be outpaced by the rate of climate change (Davis and Shaw 2001; McLachlan et al. 2005; Gray & Hamann 2013), with implications to the health and productivity of both wild forests and those planted for wood or carbon sequestration. Assisted gene flow (AGF), the “intentional translocation of individuals within a species range to facilitate adaptation to anticipated local conditions” (Aitken and Whitlock 2013), is a strategy for mitigating these deleterious effects of mismatches between genotypes and climate. For instance, warmer-adapted provenances are faster growing, although less cold hardy, for many temperate and boreal species (Aitken and Bemmels 2016; Wang et al. 2010). If genotypes are moved into suitable climates, but not so far that they suffer from cold injury or other types of maladaptation, this faster growth rate is expected to translate to higher survival, better health, and greater productivity (e.g., Wadgymar et al. 2015). When the motivation for planting is conservation, AGF could bolster the demographics of rare species or accelerate stand development for habitat and other ecosystem services. Maintaining or enhancing genetic diversity is key, as the goal of assisted gene flow in conservation settings is to establish self-sustaining populations capable of natural regeneration, establishment, and further adaptation to new conditions (Aitken and Whitlock 2013; Lunt et al. 2013; Kelly and Phillips 2015; Aitken and Bemmels 2016).

The argument for AGF with forest trees is particularly strong, due to 1) the long history of study and understanding of local adaptation to climate in many widespread species (Langlet 1971; Morgenstern 1996); 2) the lack of strong population structure and isolation that might lead to outbreeding depression (Howe et al. 2003; Neale & Savolainen 2004; Mitton & Williams 2006; Savolainen et al. 2007); 3) the long generation times of forest trees and corresponding high rate of climate change per generation (McLachlan et al. 2005; Petit and Hampe 2006; Aitken et al. 2008; Alberto et al. 2013); and 4) the infrastructure and operational practices that already exist for collecting or producing seeds, growing seedlings, and reforesting harvested or otherwise disturbed areas (Aitken and Bemmels 2016). Effective AGF strategies require an understanding of the nature of local adaptation to climate, particularly the major climatic drivers of local adaptation and how strongly populations are differentiated along these climatic gradients.

Forest scientists have traditionally used provenance trials*—in situ* field-based common garden experiments that usually involve partial reciprocal transplants—to understand links between phenotypes under divergent selection and the environments driving those differences (see discussion in Lind et al. 2018). Such designs have been the major source of knowledge of local adaptation trees for over two centuries, where differentiation among populations is usually attributed to the source environment of individuals (Langlet 1971; Morgenstern 1996). In provenance trials, phenotypic data is often limited to survival and growth rather than component traits directly related to climate, such as tolerance of cold, drought, insects, or diseases. Multi-site provenance trials can therefore provide excellent information on local adaptation, but they are limited by the decades-long time frame needed to obtain meaningful data and by the restricted geographic and climatic scopes of such trials for many species (Kawecki and Ebert 2004; Aitken et al. 2008; de Villemereuil et al 2015). Moreover, provenance trials are not feasible for some species due to a lack of available sites, sufficient resources, ethical reasons, or the difficulty of obtaining seed from many populations, particularly for endangered species or species with seed that cannot be stored (Morgenstern 1996; Blanquart et al. 2013; de Villemereuil et al. 2015; Flanagan et al. 2018).

Single or multi-environment seedling common gardens provide some advantages over traditional provenance trials. While single common gardens can be used to test for differentiation among genetic groups and to develop transfer functions (Matyas 1994; O’Neill et al. 2008), multiple common gardens can be used to test for environmental forces driving this differentiation, and need not necessarily be within source environments of populations under study. Such experiments allow for detailed phenotyping of climate-related traits at the vulnerable seedling stage that have important fitness consequences for the populations under consideration (e.g., phenology, cold-or drought-hardiness, growth, and allocation of biomass; see refs in Cornelius 1994, Howe et al. 2003, Savolainen et al. 2007; Alberto et al. 2013, and Lind et al. 2018). However, juvenile phenotypes may not reflect fitness-related traits at later life stages (e.g., reproduction) and such experimental environments are often artificial (Kawecki and Ebert 2004).

Due to the prevalence of the transplant designs mentioned above, phenotypic inferences of local adaptation and their applications to seed transfer of forest trees have been traditionally characterized in geographic terms. For example, seed transfer limits for wild-sourced seedlings in British Columbia were until recently defined as a maximum latitude, longitude, and elevation that seedling stock could be transferred from their provenance to the planting site (Ying and Yanchuk 2006). The advent of high-resolution gridded climate data over the past two decades (e.g., PRISM, Daly et al. 2002) has allowed more precise inferences of the spatial distribution of local adaptation (Wang et al. 2010) and has facilitated the transition from geography-based to climate-based seed transfer (O’Neill et al. 2017). When integrated with climate change projections (e.g., ClimateNA, Wang et al. 2016; and WorldClim, Fick and Hijmans 2017), climate data provide the essential basis for AGF and address some of the shortcomings of geographically-based (“local is best”) seed zones. While generic approaches to climate variable selection may provide a first approximation for AGF (e.g., niche modeling), information tailored to species-specific patterns relating adaptive phenotypic variation to climate will better tailor AGF strategies (e.g., as in O’Neill et al. 2017), as climatic factors limiting a species’ niche may not be those driving differentiation among populations.

In situations where phenotypic data is unavailable, genomic data could potentially be a useful alternative to phenotypic data for inferring the climatic drivers of local adaptation. Population genomic approaches for detecting adaptive variation have become feasible within the last decade (Neale & Savolainen 2004; Sork et al. 2013; Prunier et al. 2015; Lind et al. 2018). Next generation sequencing methods now allow for the genotyping of large numbers of variants (e.g., single nucleotide polymorphisms, SNPs) in non-model species for elucidating aspects of the species biology that can inform management and conservation decisions (Lotterhos et al. 2018; Mähler et al. 2017; Flanagan et al. 2018; Parchman et al. 2018; Rellstab et al. 2018). Genotype-environment association (GEA) approaches can identify both the environmental drivers of local adaptation and loci underlying locally adaptive traits (Schoville 2012; De Mita et al. 2013; Rellstab et al. 2015). Likewise, genotype-phenotype association (GPA) studies can identify loci associated with climate-related phenotypes (Neale & Savolainen 2004; Prunier et al. 2015; Holliday et al. 2017). These methods can be combined to identify suites of potentially locally adapted loci (e.g., Yeaman et al. 2016; references in Lind et al. 2018). Despite the extensive literature on the genomic basis of local adaptation, however, we are not aware of any operational uses of genomic data to guide seed transfer or AGF.

Genomic data have many potential roles in guiding AGF as an alternative or a supplement to phenotypic and climatic data. Given the unavoidable costs and lag time of provenance trials and common garden experiments, the prospect of characterizing local adaptation using genomics rather than phenotypes is appealing. Genomic approaches could bring down the cost and response time of managing AGF for commercially important tree species, and also provide the opportunity for comprehensive (or strategic) genetic conservation of many species that lack resources for phenotypic trials. In addition to being a potential alternative to phenotypes, genomic data can provide unique insights into local adaptation that are not available from phenotypic or climatic data alone. For example, rangewide phenotypic clines can potentially mask more localized allelic clines that underlie adaptive traits (see Box 1). Similarly, the spatial structure of standing variation in adaptive alleles—an important consideration for AGF and *in situ* genetic conservation—can only be inferred from genomic data.

The objective of this study is to evaluate genomic data, relative to phenotypic and climatic data, as a basis for assisted gene flow and genetic conservation of locally-adapted conifers. We address three research questions using phenotypic and genomic data from 281 provenances of *Pinus contorta* Dougl. ex Loud from across western Canada. Firstly, *what is the relative value of genomic data vs. climatic and geographic data in explaining locally adaptive phenotypic variation?* We address this by comparing the proportion of variance in four seedling traits that can be explained by geographic, climatic, and several types of genomic data including a full SNP array, a large set of neutral markers, and loci inferred from both genotype-phenotype associations and genotype-environment associations. Secondly, *can genomic data identify the climatic drivers of local adaptation?* We use phenotypic data from both a short-term common garden study and a long-term provenance trial to contrast the predicted importance of various climatic drivers of phenotypic differentiation to that predicted from genomic data (GEA loci). Thirdly, we examine information that is uniquely available from genomic data—the genetic clines underlying phenotypic clines—to address the question: *what is the spatial scale of local adaptation to climate?*

These assessments identify the contributions that genomic data can make to assisted gene flow and genetic conservation in a changing climate.

#### BOX 1 The structure of allelic variation underlying phenotypic clines in adaptive traits

For widespread tree species that experience both strong diversifying selection and high gene flow, climatic gradients often drive clinal variation in phenotypes (Endler 1977; Alberto et al. 2013). However, the number and geographic distribution of adaptive loci underlying these patterns is, for the most part, unknown.

There are two ways for genetic clines to produce a rangewide cline in an additive polygenic trait (Figure 1). The first is to have concordant clines in the underlying loci, representing a gradual rangewide shift in allelic frequency across all underlying loci (Figure 1B) that therefore matches the range-wide phenotypic cline (Figure 1A). Alternatively, a phenotypic cline can result from multiple distinct, localized genetic clines, each providing variation sequentially over short sections of the environmental gradient (Barton 1999; see also Box 3 in Savolainen et al. 2007), as depicted in Figure 1C.

The degree to which local adaptation is structured as localized, sequential genetic clines has implications for AGF, as this may reduce the amount of standing adaptive variation and thus adaptive potential. Ultimately, the spatial scale of adaptation is a function of gene flow, selection, and drift. In species with long-isolated populations and little gene flow, such structure could also risk lower compatibility between native and transplanted individuals, but outbreeding depression is unlikely in widespread, abundant, wind-pollinated trees (Aitken and Whitlock 2013). If adaptive variation is distributed as concordant range-wide genetic clines, loci underlying an adaptive trait will be polymorphic throughout most of the species range, except perhaps at the range margins, or in otherwise isolated or small populations. In this case, standing variation should exist for adaptive loci that could enable *in situ* adaptation to climatic change, as long as locally novel climatic conditions exist elsewhere in the species range and are not isolated from gene flow. Localized clines, in contrast, imply that standing variation in a subset of adaptive alleles is limited to only a portion of the species’ range.

**Figure 1:**
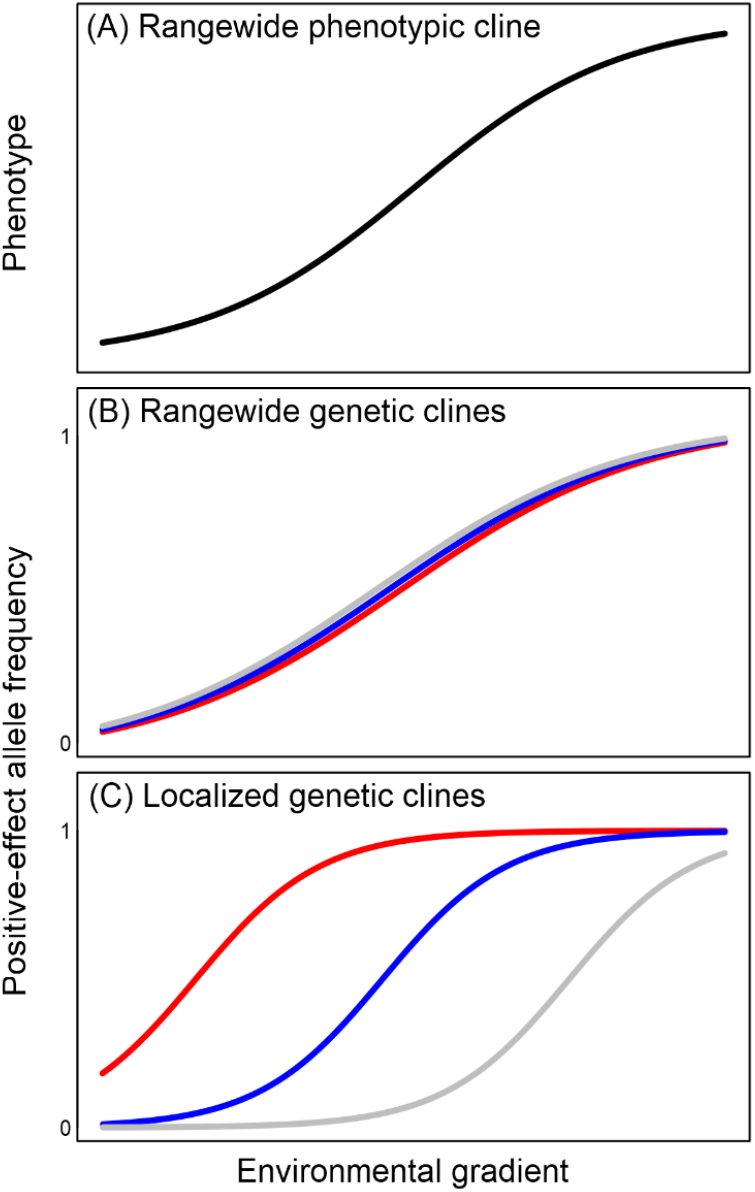
Illustration of rangewide vs. sequential, localized genetic clines (B-C) underlying a continuous phenotypic cline (A) along an environmental gradient (After Barton 1999).

## 2 Methods

### 2.1 Phenotypic data

#### 2.1.1 Seedling common garden experiment

The primary phenotypic data in this study originate from a raised bed common garden of 1,594 lodgepole pine seedlings at Totem Field at the University of British Columbia in Vancouver, BC. Design, establishment, and measurement of the common garden, summarized here, are described in detail by MacLachlan et al. (2017). Briefly, seedlots originated from 281 provenances representing lodgepole pine’s climatic range within British Columbia and Alberta (Figure 2E). Seedlots were predominantly selected from the range of the Rocky Mountain subspecies (*P. contorta* Dougl. ex Loud. ssp. *latifolia* [Engelm.] Critchfield), but also include the coastal subspecies (*P. contorta* Dougl. ex Loud. ssp. *contorta)* and the region of hybridization with jack pine (*Pinus banksiana* Lamb.) in northern Alberta.

**FIGURE 2.**
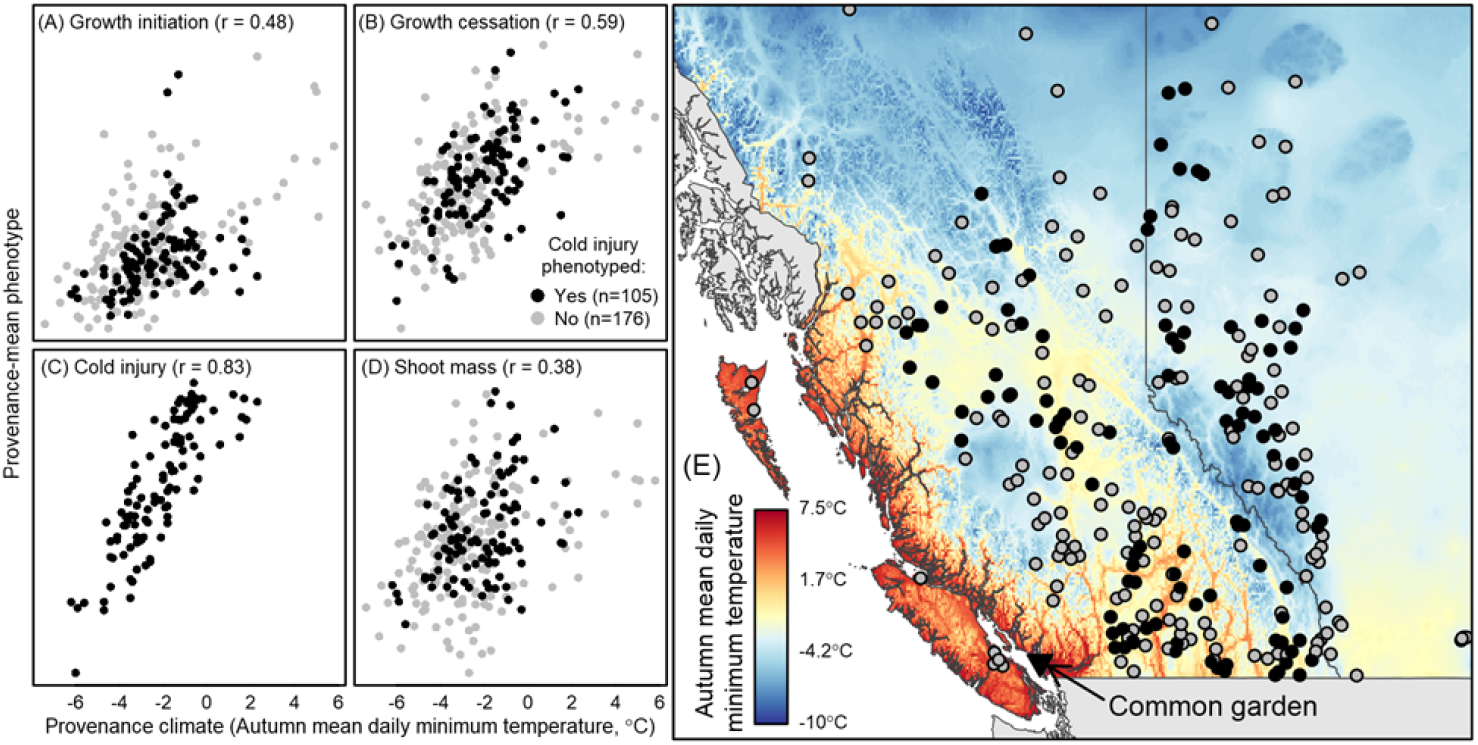
Phenotypic clines of four traits in lodgepole pine seedlings grown in a common garden. A total of 1,594 seedlings from 281 provenances across British Columbia and Alberta, Canada (grey and black circles) were phenotyped for growth initiation (A), growth cessation (B), and three-year shoot mass (D). A subset of 922 seedlings from 105 provenances (black circles) were tested for autumn cold injury (C). Phenotypic clines (A-D) are plotted on an environmental gradient of autumn mean daily minimum temperature, mapped in (E).

Our study utilizes phenotypic data from four traits: growth initiation, growth cessation, autumn cold injury, and shoot mass (methods in MacLachlan et al. 2017). We removed experimental effects from phenotypic values by reporting phenotypes as *z*-standardized residuals of a linear mixed effects model, implemented with ASreml-R (Butler 2009), in which experimental block and location within block are random effects:

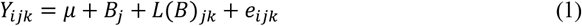

where *Y_ijk_* is the phenotypic observation of a trait made on individual *i* grown in the *j*^th^ block (5), at the *k*^th^ seedling location (*L*) nested within block (*L*(*B*)_jk_), *μ* is the experimental mean, and *e* is the residual error of individual *i*.

#### 2.1.2 Illingworth provenance trial

We analyzed 20-year heights from the Illingworth lodgepole pine provenance trial to corroborate the inferences from the Vancouver seedling common garden with longer-term data from sites more typical for this species. This trial, established in 1974 by the BC Ministry of Forests (Illingworth 1978; Wang et al. 2010), tested a rangewide (New Mexico to Yukon) collection of 140 provenances at 60 sites in interior British Columbia. We assessed the strength of the univariate relationships between 20-year height and 19 climate variables for three contrasting trial sites: one each from southern (PETI), central (NILK) and northern (WATS) British Columbia (Supp. Info Figure S1). An adjusted *R*^2^ was estimated for the quadratic relationship between provenance climate and the average 20-year heights of the provenances at each test site. This relationship was estimated for each of the 19 standard climate variables (Table 1) used in this study. Reported results are the mean *R*^2^ over the three sites.

### 2.2 Climate data

Climate normals 1961-1990 period for each provenance in the seedling common garden were obtained from ClimateNA (Wang et al. 2016), using the latitude, longitude, and elevation of each seedlot. The 19 bioclimatic variables used in this study (Table 1) are the same as used in previous analyses of genomic datasets from the AdapTree Project, selected *a priori* based on relevance to the species biology and environmental variation across provenances (Yeaman et al. 2016a; MacLachlan et al. 2017; Lotterhos et al. 2018). In addition to these 19 analysis variables, we use autumn mean daily minimum temperature (Tmin_at) as the environmental gradient for plotting phenotypic and genetic clines. We selected Tmin_at due to its biological relevance to autumn cold injury, growth cessation, and shoot mass.

**TABLE 1.** The set of 19 bioclimatic variables used in this study

| Environmental Variable (unit) | Abbreviation |
| --- | --- |
| Mean annual temperature (°C) | MAT |
| Mean warmest month temperature (°C) | MWMT |
| Mean coldest month temperature (°C) | MCMT |
| Continentality (MWMT minus MCMT) (°C) | TD |
| Mean annual precipitation (mm) | MAP |
| May to September precipitation (mm) | MSP |
| Annual heat-moisture index (MAT+10)/(MAP/1000) (°C/μm) | AHM |
| Summer heat-moisture index ((MWMT)/(MSP/1000)) (°C/μm) | SHM |
| Degree-days below 0°C, chilling degree-days | DD_0 |
| Degree-days above 5°C, growing degree-days | DD_5 |
| Number of frost-free days (days) | NFFD |
| Frost-free period (days) | FFP |
| The day of the year on which FFP begins (Julian date) | bFFP |
| The day of the year on which FFP ends (Julian date) | eFFP |
| Precipitation as snow between August and July (mm) | PAS |
| Extreme minimum temperature over 30 years (°C) | EMT |
| Extreme maximum temperature over 30 years (°C) | EXT |
| Hargreaves reference evaporation (mm) | Eref |
| Hargreaves climatic moisture deficit (mm) | CMD |

### 2.3 Genomic data

#### 2.3.1 SNP table

DNA was extracted from tissue of spring needles using a Macherey-Nagel Nucleospin 96 Plant II Core™ kit, automated on an Eppendorf EpMotion 5075™ liquid handling platform. Samples were genotyped by Neogen GeneSeek (Lincoln, Nebraska) using the AdapTree lodgepole pine Affymetrix Axiom 50K lodgepole pine SNP array. SNP discovery for this array was based on the lodgepole pine sequence capture dataset described by Yeaman et al. (2016) and Suren et al. (2016). It included probes for the exons of 24,388 genes, as well as intergenic regions, with intron-exon boundaries identified by mapping the lodgepole pine transcriptome to the loblolly pine (*Pinus taeda* L.) v1.01 draft genome (Neale et al. 2014, Zimin et al. 2014). SNPs were selected for inclusion based on preliminary GEA analyses as well as GPA using phenotypes for seedling traits (Yeaman et al. 2016), differentially-expressed genes (Yeaman et al. 2014), candidate genes for climate adaptation from other conifers, mappable SNPs for a linkage map, and a set of non-coding loci to control for neutral population structure. Genotypes from the SNP table were filtered to retain 36,384 SNPs with a minor allele frequency ≥ 0.01. Of these filtered loci, 4,750 selectively neutral SNPs (those intergenic SNPs that had no significant genotype-phenotype or genotype-environment associations in the analyses of Yeaman et al. 2016) were selected for population structure correction in association analyses. Excluding this “neutral set”, the final candidate adaptive SNP table used in associations contained 31,634 SNPs. We genotyped 1,594 seedlings from the Vancouver outdoor common garden and an additional 1,906 seedlings from the same 281 provenances grown in a separate growth chamber experiment (Liepe et al. 2016), for a total median sample size of 11 seedlings (range seven to 24) for each provenance (Figure S2).

#### 2.3.2 Genotype-Phenotype Association (GPA)

We implemented GPA using the phenotypic residual values (from Eq. 1) for each of the four traits measured at the Vancouver common garden using the linear regression-based *mlma* function in GCTA (Yang et al. 2011). We corrected for population structure using the *grm* option of *mlma* with the 4,750 putatively neutral SNPs described in *§2.3.1.* We limited marker data to one SNP per contig to reduce redundancies due to physical linkage, which reduced the number of available SNPs from 31,634 to ~19,600 SNPs. SNPs in the bottom 1% of GPA p-values for each trait were identified as candidate SNPs (*n* = 196 SNPs per trait). For each candidate SNP, the allele that increased the value of a phenotype – called the positive effect allele (PEA) – was identified from the regression slope in the GCTA *mlma* output.

#### 2.3.3 Genotype-Environment Associations (GEA)

We used bayenv2 (Coop et al. 2010; Günther & Coop 2013) to identify loci with evidence for responses to environmental selection. The neutral covariance matrix for this analysis was estimated using the set of 4750 neutral loci for 100,000 iterations. For each centered and standardized environmental variable (Table 1), we ran bayenv2 in test mode for one million iterations across three independent chains for the 31,634 loci that did not overlap with the neutral set, using the covariance matrix to correct for neutral population structure. To reduce the marker set to one SNP per contig (~19,600 SNPs), we retained loci that had the greatest evidence for environmental response from each contig (average rank across absolute *rho* and Bayes factor [BF] across the three chains; i.e., six values). To ensure we isolated only loci with the strongest evidence for environmental influence, we re-ranked these ~19,000 loci and retained only those that met two criteria for a given environmental association: 1) the locus was in the top 300 ranked loci for BF for each of the three chains, and 2) was also in the top 300 ranked loci for absolute value of *rho* for each of the three chains. In addition to using these GEA loci towards our objectives, we report the number of loci identified using these criteria, as well as the overlap between GPA and GEA.

### 2.4 Analyses

We present three analyses that correspond to the three research questions posed in the final paragraph of the Introduction.

#### 2.4.1 Phenotypic variation explained by geographical, climatic, and genomic data

One way of assessing the relative value of geographic, climatic and genomic data for guiding assisted gene flow and other climate adaptation strategies is to measure the degree to which they can be used to statistically explain locally adaptive phenotypic variation. The dimensionality of the information in each data source is expected to differ: for example, genome-wide data may be distributed over many more modes of variation than the three dimensions (latitude, longitude, elevation) required to fully describe geographic location. These data sources can compared on equal terms by extracting their principal components (PCs) and assessing the cumulative explanatory content of increasing numbers of PCs as predictor variables. Explanatory content in this case is measured as proportion of variance explained (*R*^2^) by a multivariate regression of phenotypic values (the response variable) against the PCs of the geographical, climatic, or genomic data (the predictor variables). We used multiple linear regression for this purpose, and report the mean *R^2^* of a 5-fold cross-validation implemented with the *cv.lm* function of the DAAG package in R (R Core Team 2017). For comparison, we also performed this analysis with Random Forest regression, a regression tree ensemble learning algorithm that provides cross-validated modeling of non-linear relationships and variable interactions (Breiman 2001). For this analysis, we selected a subset of *climate-associated* GPA loci with *R*^2^>0.2 in multiple linear regressions on the 19 climate variables specified in Table 1.

#### 2.4.2 Climatic drivers of local adaptation

We examine the congruence of genomic vs. phenotypic data in guiding climatic variable selection by contrasting the proportion of variance of individual climate variables that is explained by climate-associated genomic loci, seedling common garden phenotypes, and long-term provenance trial phenotypes. For each data source, we conducted one regression for each of the 19 climate variables (Table 1), in which the response (dependent) variable is the provenance climates for a single climate variable. The predictor (independent) variables for the genomic regressions are the first four principal components of the minor allele frequencies for the top-300 GEA loci associated with the climate variable of interest (see *§2.3.3* for GEA methods). The predictor variables for the seedling common garden regressions are the provenance means of the standardized phenotypes for the four traits (see *§2.1.1).* The predictor variables for the long-term provenance trial are the 20-year heights measured at three sites of the Illingworth trial (see *§2.1.2).* Note that the Illingworth data sample a different set of provenances than the genomic and seedling common garden data, and thus are essentially independent of these two other data sources. As in the previous analysis (*§2.4.1*), we used multiple linear regression and report the mean *R*^2^ of a five-fold cross-validation for each regression.

#### 2.4.3 Spatial scale of local adaptation to climate

To characterize the genetic clines associated with the seedling traits measured in the common garden, GPA loci were clustered using a Euclidean k-means algorithm (*kmeans{*stats}; R Core Team 2017). To cluster SNPs, we transposed the provenance-mean positive-effect allele frequency data so that SNPs occupied the row (observations) position and provenances occupied the column (variable) position. Clusters, then, are SNPs that have similar allele frequencies across provenances. Similarity in this configuration is distinct from correlation: SNPs with large differences in aggregate allele frequency will be put in separate clusters, even if they are very highly correlated. Hence this clustering approach is distinct from standard LD clustering approaches based on allele frequency covariance. We use the cluster mean positive-effect allele frequency for each provenance to visually summarize the clusters. Averaging reduces variance, however, which distorts genetic clines. To restore the variance of the cluster mean positive effect allele (PEA) frequency, we multiplied the cluster-mean PEA frequency for each provenance by the mean standard deviation of the SNPs in the cluster.

To investigate levels of standing variation, we calculated expected heterozygosity (*H*_e_) for each PEA in each provenance. The cluster-mean *H*_e_ for each provenance is the mean *H*_e_ for each SNP within the cluster. We report standing variation as proportional polymorphism for each provenance: the proportion of SNPs within a cluster with *He* > 0.

## 3 Results

### 3.1 Phenotypic clines

Provenance-mean phenotypes for all four traits measured in the Vancouver common garden exhibit moderate to strong clines relative to study area temperature gradients where the timing of growth cessation and fall cold injury show the strongest relationships with many climate variables, such as autumn mean daily minimum temperature (Figure 2). In general, trees from colder provenances initiated growth slightly earlier, ceased growth earlier, achieved less total growth, and exhibited less cold injury. Autumn cold injury in particular has a very strong relationship (r = 0.83) to autumn temperature. Within-provenance variation among individuals is generally uncorrelated among the four traits (Figure S3). However, within-provenance variation of shoot mass is positively correlated to growth cessation day (*r*=0.59) and weakly but significantly negatively correlated to growth initiation day (*r* = −0.18, *p* = 2_E_ −12). This result may be due to the benign maritime climate of the common garden; seedlings with a longer growth period are not penalized by environmental constraints such as growing season frosts. Correlations of among-provenance variation in growth cessation, fall cold injury, and shoot mass are moderate to strong. Growth initiation is poorly correlated with the other traits.

### 3.2 Phenotypic variation explained by geographical, climatic, and genomic data

The absolute and relative explanatory content of geographical, climatic, and genomic data differs among traits (Figure 3). Differences in the explained phenotypic variation among traits generally exceed the differences among the three types of data (geographic, climatic and genomic) within traits, and mirror the strength of the phenotypic clines in Figure 2. Nevertheless, there are important differences in the relative explanatory content of geographic, climatic, and genomic data among traits. In general, geographic variables (yellow diamonds) are as predictive of seedling phenotypes as climatic variables (gray circles, Figure 3), consistent with strong local adaptation to geographically-based climate in this species. The exception is growth initiation, where geographic variables are more explanatory than climate. The GPA SNPs (solid black line, Figure 3) are more explanatory than climate and geography in growth initiation and shoot mass but not growth cessation, where they are equivalent, and cold injury, where they are slightly inferior.

**FIGURE 3.**
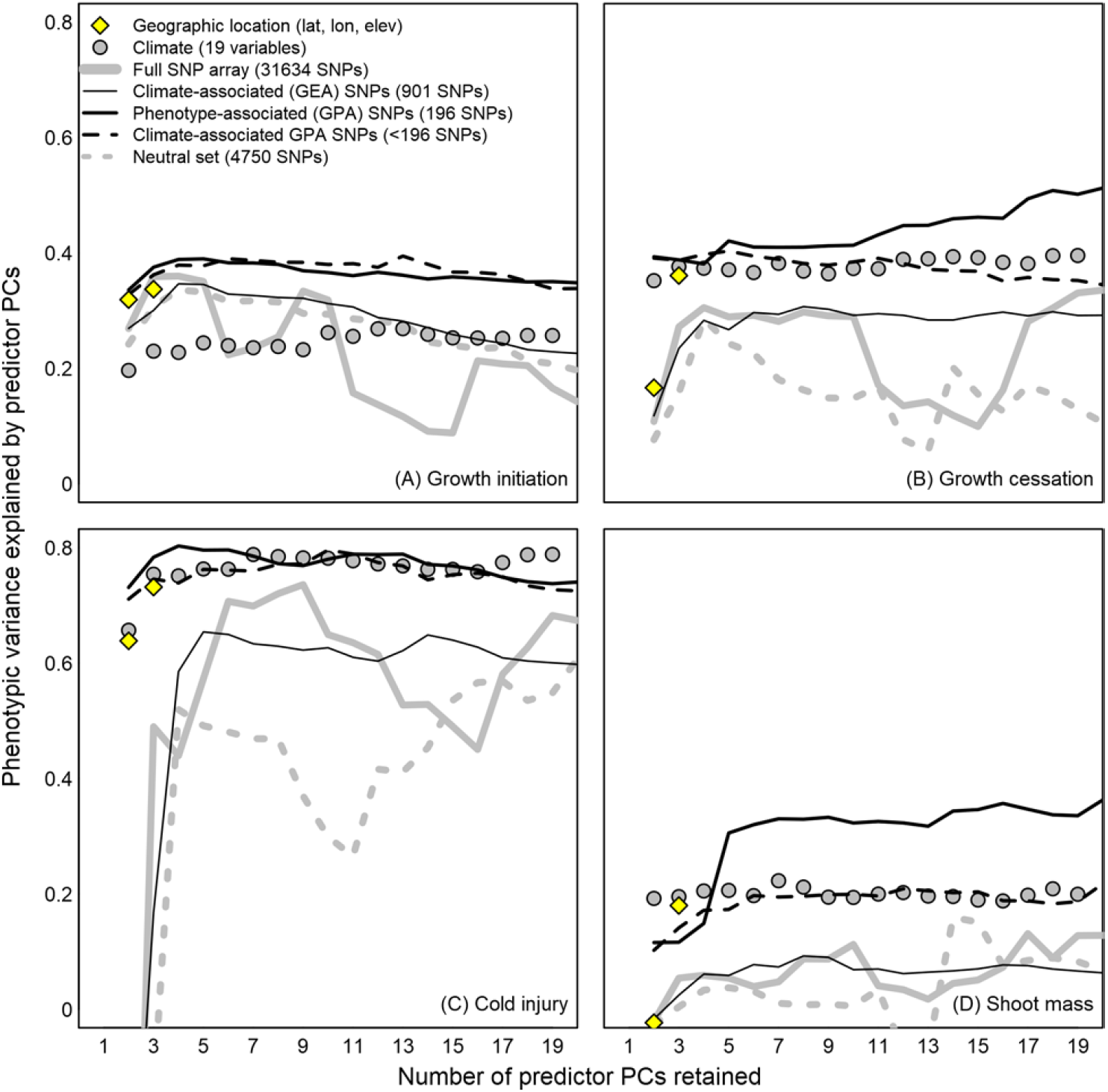
Common garden phenotypic variance explained (PVE) for four traits by cumulative principal components of geography (diamonds), climate (circles), and several subsets of genomic data from a SNP array (lines). Each point is the cross-validated *R*^2^ of a multiple linear regression of provenance-mean phenotype against the specified number of principal components of the predictor data. GEA SNPs (thin black line) are the pooled top-300 SNPs based on Bayes factor from each of the 19 climate variables. GPA SNPs (thick black line) are the top 1% of coding-region SNPs (maximum of one SNP per contig) based on the p-value of a population-structure-corrected linear association of allele frequencies to seedling phenotypes. Climate-associated GPA SNPs (black dashed line) are GPA SNPs with a linear association to climate (see *§2.4.1*). The neutral set is shown as a grey dashed line.

The relative explanatory power of different types of genomic data is consistent among traits (Figure 3), and provides several insights. First, GPA SNPs (solid black line) consistently have the highest explanatory power. Since the GPA SNPs are a subset of the full array (solid gray line), the difference between GPA and full SNP array indicates the value of extracting the relevant genetic information. Second, the climate-associated GPA SNPs (black-dashed line, Figure 3) generally explain less phenotypic variation than the full set of GPA SNPs. In the case of growth initiation, however, climate-associated GPA SNPs explain more phenotypic variation than climate variables. Third, the GEAs identified using bayenv2 (Supplemental Table S1) consistently have low explanatory power to predict phenotypic variation, but higher and more stable explanatory power than the neutral set and the full SNP array. There is a fairly high overlap of GEA with GPA loci, with an average of 53% of GEA SNPs from various environmental variables found within 1000 bp of GPA loci (range 0% for NFFD to 81% for EXT; sd = 20.6%), and a total of 35% of GPA SNPs found within 1000bp of the GEA loci found across environmental variables (Supplemental Table S1). This is another line of evidence of the strong role of climate in driving phenotypic variation among provenances.

The neutral set and full SNP array both have explanatory relationships with phenotypes, but these are not as strong as relationships with geographic, climatic, and filtered genomic data (Figure 3). An equivalent analysis to Figure 3 using Random Forest regression instead of linear regression demonstrates that both the neutral set and full SNP array contain almost as much non-linear explanatory information as the climatic and geographic variables (Figure S4). Further, some subsets of the neutral set exhibit linear relationships to phenotype that are as strong and stable as the relationships of GEA loci to phenotype (Figure S5).

Traits differ substantially in the dimensionality of their associated genomic information, i.e., the number of PCs at which further gains in explanatory information are not achieved. Explainable variation in growth initiation, growth cessation, and autumn cold injury are almost completely described by the first two PCs (Figure 3). In contrast, five PCs are required to describe the explainable variation in shoot mass. The dimensionality of explanatory information in the different traits speaks to the complexity of genetic controls on the trait.

### 3.3 Climatic drivers of local adaptation

The GEA loci show general congruence with both the short-term (3-yr) common garden experiment (Figure 4A) and a longer-term (20-yr) provenance trial (Figure 4B). Across both phenotypic traits and the genomic GEA data, there is agreement that local adaptation is strongly associated with winter temperature variables: mean temperature of the coldest month (MCMT), degree-days below 0°C (DD_0), winter-summer temperature contrast (TD), and extreme minimum temperature (EMT; variables in upper right of Figure 4A and 4B). Note that mean annual temperature can be considered primarily a winter variable in this study area because spatial variation in mean temperature along the latitudinal gradient is much stronger in winter than in other seasons. In the Vancouver common garden (Figure 4A), this congruence between genotypic and phenotypic relationships to climate variables is broken by summer temperature variables (Eref, EXT, DD5, and MWMT), which have moderate associations with phenotypes (x-axis) but low associations with genotypes (y-axis). In the provenance trial (Figure 4B), the congruence is broken by summer precipitation variables (MSP and CMD), which have low associations with phenotype but moderate associations with genotype. The same pattern of these relationships is produced using either the full SNP array or the neutral SNPs in place of the GEA SNPs (Figures S6 and S7, respectively).

**FIGURE 4.**
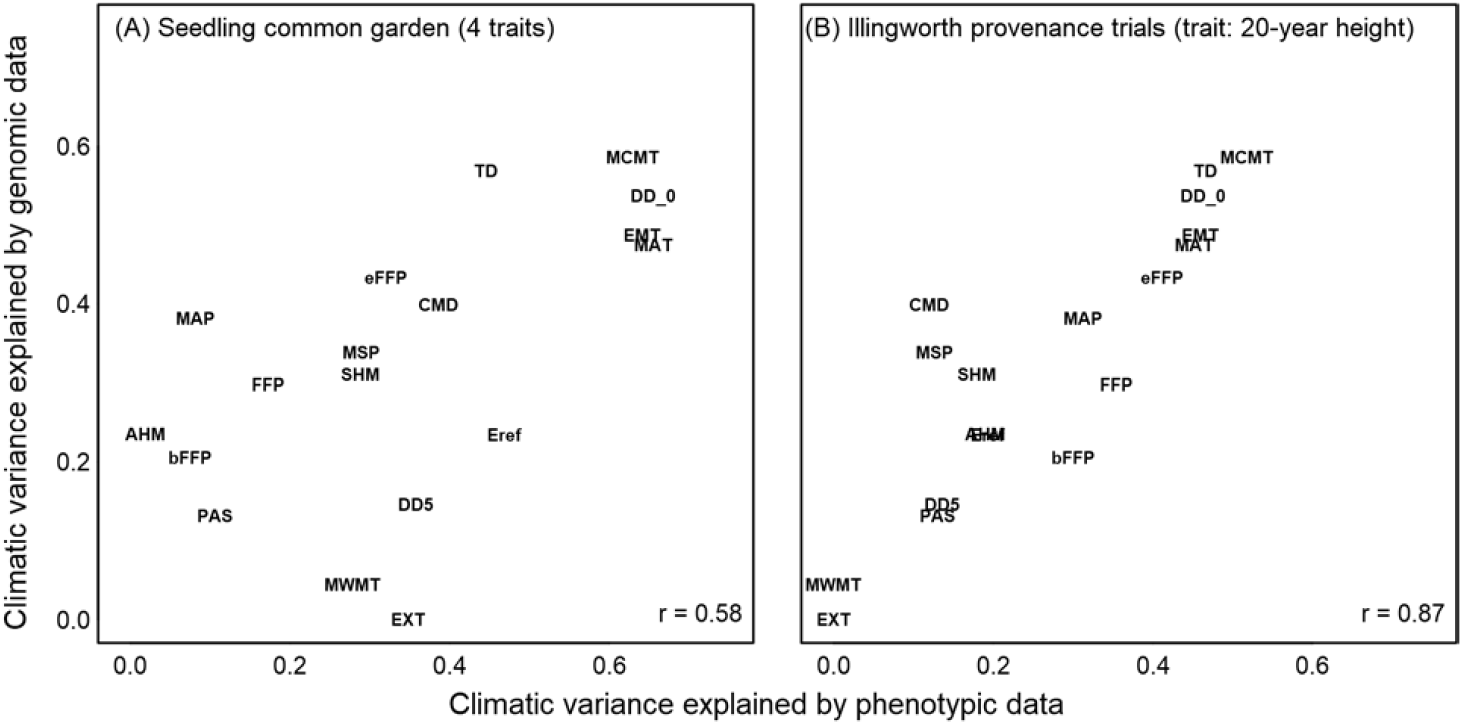
Climatic variable selection based on phenotypic vs genomic data. Variance explained is the crossvalidated *R*^2^ of a multiple linear regression of each climate variable (response variable) against the phenotypic or genomic predictor variable set. Genomic data (predictor variables for the y-axis analyses) are four principal components of the minor allele frequencies for the top-300 GEA SNPs identified by bayenv2 for each climate variable. Phenotypic data (predictor variables for the x-axis analyses) for panel A are provenance-mean phenotypes for the four common-garden traits presented in Figure 2. Phenotypic predictor data for panel B are 20-year heights of the Illingworth lodgepole pine provenance trial. Climate variable acronyms are described in Table 1.

### 3.4 Spatial scale of local adaptation to climate

All four common garden traits exhibit linear phenotypic clines over many of the climatic gradients of the study area, where the strongest of these clines is autumn cold injury relative to autumn temperature (Figure 2C; *r* = 0.83). To detect whether genetic clines for cold injury loci along environmental gradients are rangewide or localized, we examined the *n* = 80-locus subset of the 196 cold injury GPA candidates that are also moderately associated with the 19 climate variables (Random Forest pseudo-*R*^2^ > 0.31; Figure S8). We clustered these 80 loci into six clusters based on their absolute PEA frequencies across provenances (Figure S9). The within-cluster mean PEA frequencies of these six clusters have distinct clines (Figure 5) relative to the gradient in autumn temperature across the study area (Figure 2). Clusters 2, 4, and 5 show no clinal variation across provenance temperatures below −3°C, but have a clinal increase in PEA frequency across higher temperatures (Figure 5). Cluster 6 has essentially the opposite pattern, in that it shows clinal variation almost exclusively below the −3°C autumn temperature threshold. The adaptive variation in cluster 6 is of particular interest, in the context of standing variation, because it is localized to a high degree relative to the other clusters. Cluster 1 has an inverse pattern to cluster 6 relative to provenance climate, and primarily reflects variation associated with the coastal ssp. *contorta*, which occur at Tmin_at >2°C. Cluster 3 exhibits increased variation in the interior of BC, which appears to be reversed in the warmer climates of the coast.

**FIGURE 5.**
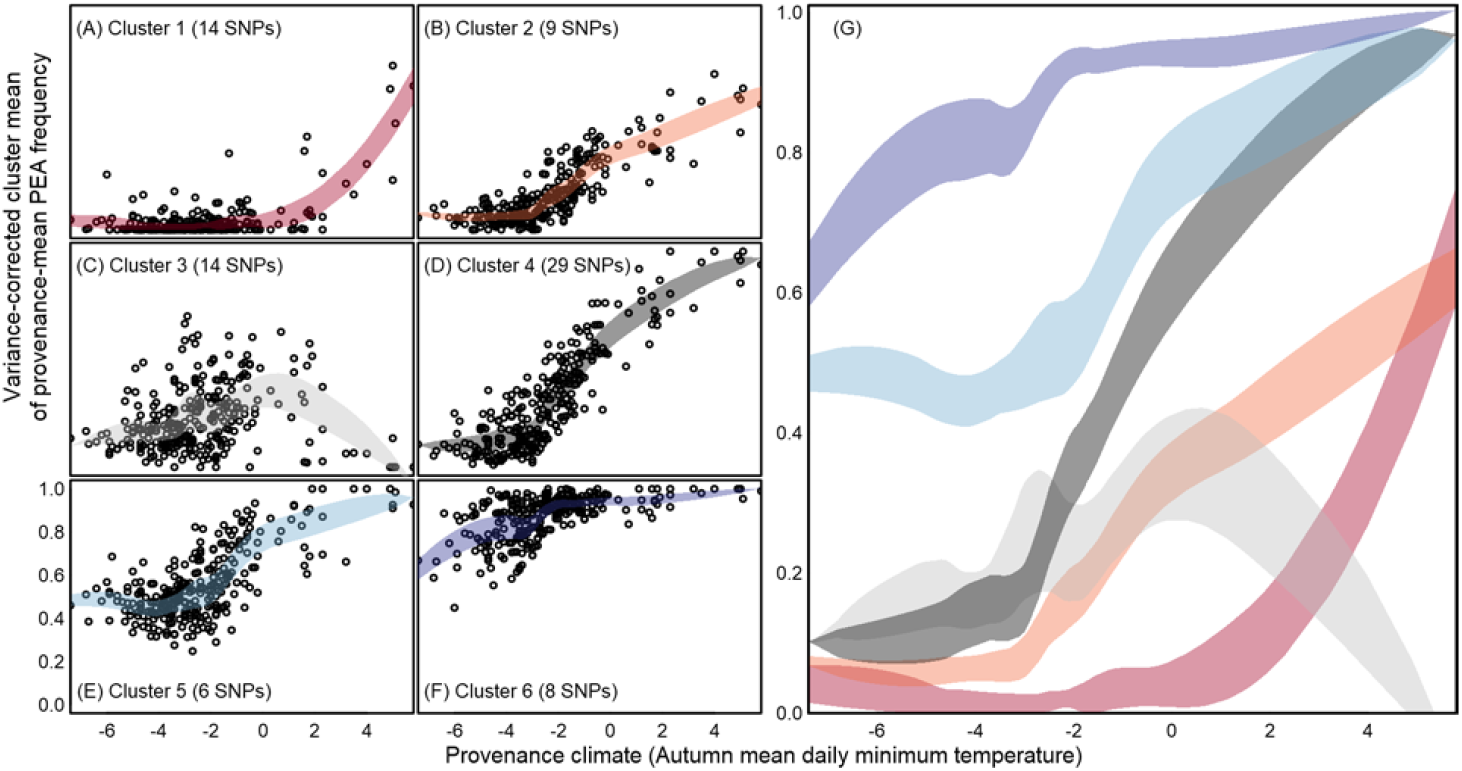
Genetic clines associated with autumn cold injury. (A-F) the 80 climate-associated GWAS SNPs for autumn cold injury are clustered based on similarities in positive effect allele (PEA) frequencies across provenances (n=281). Each point is the mean of the PEA frequencies across clustered SNPs for one provenance, with a correction applied to restore the variance of the PEA frequencies following averaging. The colored bands in each plot, superimposed in panel G, are locally-weighted 0.5-standard deviation prediction intervals. Recall that the y-axes are reflecting the frequency of PEAs that are associated with increased cold injury

To contrast the extent of rangewide vs localized clines, the geographic patterns of allele frequencies in clusters 4 and 6 are shown in Figure 6. Cluster 4 represents the dominant rangewide genetic cline over the study area, and is largely parallel with clusters 2 and 5. Cluster 6 is the complementary cline to cluster 4 as it reflects adaptive variation for cold hardiness in boreal provenances. Cluster 4 has a strong cline with respect to the joint thermal gradient of latitude and elevation (Figure 6C). Putatively adaptive alleles of cluster 6 are predominantly found in the Boreal climates of Northern Alberta, Northeastern BC, and the eastern foothills of the Rocky Mountains (Figure 6F). Unlike cluster 4, cluster 6 does not have a pronounced elevational cline at low latitudes (Figure 6D). With the exception of two coastal provenances, all provenances have standing variation in some of the adaptive alleles in each cluster, though several provenances west of the Rocky Mountains (i.e., in British Columbia) have no standing variation in at least half of the cluster 6 loci (Figure S10).

**FIGURE 6.**
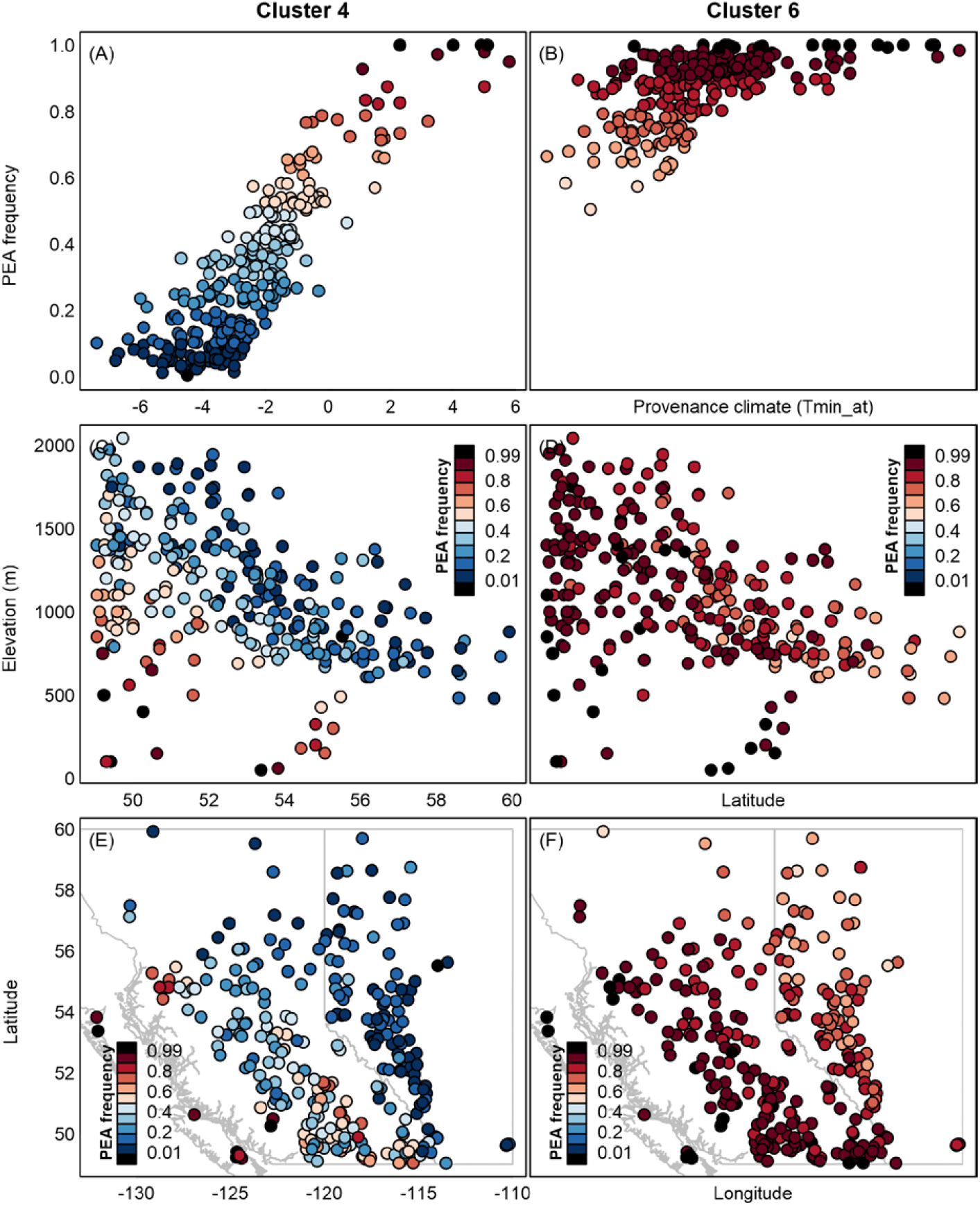
Contrasting geographic patterns of standing variation in rangewide and localized genetic clines associated with autumn cold injury. A rangewide cline (Cluster 4, left column) and a localized cline (Cluster 6, right column) relative to the autumn temperature gradient (Tmin_at) in the sampled provenances (A and B, respectively) as previously shown in Figure 4D & 4F. These clines are also compared across latitude and elevation (C,D), and latitude and longitude (E,F). Populations are colored with respect to PEA frequency (alleles that are associated with an increase in autumn cold injury).

## 4 Discussion

This study uses a large sample of locally adapted *P. contorta* provenances from across western Canada to evaluate genomic data, relative to phenotypic and climatic data, as a basis for assisted gene flow and genetic conservation. In the introduction, we posed three research questions related to this objective. The first was: *what is the relative value of genomic data vs. climatic and geographic data in explaining variation in locally adapted traits?* All data types identified the importance of adaptation to seasonally low temperatures driving population differentiation across western Canada. On average, the best predictors of seedling traits were the GPA SNPs, both the full set and the subset that was also associated with climate (Figure 3). For cold injury and growth cessation, the climate variables had similar predictive power to the GPA SNPs. For all traits, the neutral and GEA SNPs explained far less variation than climate, GPA SNPs or even geographic coordinates. This suggests that loci associated with climate-relevant traits within populations are both effective for revealing adaptive differences among populations, as well as elements of genetic architecture underlying adaptive responses that can be useful guiding management or conservation decisions.

The second question was: *can genomic data identify the climatic drivers of local adaptation?* Genotype-environment associations and a long-term provenance trial had strong agreement on the climatic drivers of local adaptation, namely winter temperature-related variables (Figure 4). These dominant drivers also held for short-term common garden seedling traits, though the overall relationship to the GEA result was weaker. This suggests that genomic data can be a viable option for identifying the key climatic controls on productivity and lifetime fitness, and may even be more reliable for this purpose than seedling traits in some contexts (Figure S11).

The third question was: *what is the spatial scale of local adaptation in climatically adaptive traits?* We found that some of the genetic clines associated with the observed phenotypic cline in cold injury are constrained to the extremes of the study area (Figure 5). However, we did not find compelling evidence for highly localized genetic clines at scales that would constrain local seed transfer to scales finer than those indicated by phenotypic data or necessitate geographically small genetic conservation units (Figures 5 and 6).

### 4.1 Phenotypic variation explained by geographical, climatic, and genomic data

The predictive power of climate variables, geography, and genotypes varied greatly among seedling traits. It is widely recognized that cold hardiness shows strong population differentiation in most temperate and boreal species (Howe et al. 2003; Alberto et al. 2013; Aitken and Bemmels 2016), and we found strong population differentiation for cold injury, as well as high predictability of cold injury from climatic, geographic, and GPA SNP data (PVE > 0.6). However, the remaining traits were not strongly predicted with any of the given data (PVE < 0.5, Figure 3). Variability in the predictive ability among traits for a given data source, or among data sources for a given trait may be due to several factors, including (discussed in Lind et al. 2018): 1) how well each phenotype is correlated with lifetime fitness; 2) the degree to which the trait is polygenic; 3) the mode of gene action underlying the genetic architecture of the trait (e.g., additive, epistatic/GxE, or pleiotropic); 4) the primary source of genetic variation in a trait (i.e., protein coding or regulatory regions); 5) the degree to which selection has structured variation within the species (i.e., the joint effects of selective forces and demographic dynamics); or 6) shortcomings of methodologies (e.g., correcting for population structure that could remove adaptive signals that covary with demography).

While this study focussed on relatively few seedling traits, there are undoubtedly many other traits at various life history stages that have population differences associated with local climate (e.g., biotic and abiotic responses, reproduction, and tree form). Our GPAs specifically identify SNPs associated with our focal seedling traits, and so it is not surprising that the GPA SNPs from individual seedling traits were better predictors of a given trait than the GEA SNPs (Figure 3). Even so, the GPA SNPs were consistently the best set of markers for explaining variation in phenotypes, and second only to climate for growth cessation and cold injury (Figure 3), emphasizing the added value of these candidate loci. Climate also consistently explained phenotypic variation well, relative to genomic data, for traits other than growth initiation. Geographic coordinates (latitude, longitude, and elevation) predicted all seedling traits quite well, explaining the success found in the vast body of older genecological literature in forest trees that used geographic variables as a proxy for climate before spatial climatic data became widely available.

In line with expectations of polygenic architectures for most of the traits, the entire SNP array (~31K SNPs) was able to predict some of the variation in these traits. Neutral SNPs from non-coding regions of the genome were also able to explain a substantial portion of phenotypic variation in all traits except shoot mass (Figure 3), and were equivalent to all other data sources as a predictor set for Random Forest regressions (Figure S4). The predictive power of neutral SNPs emphasizes the potential to confound neutral population structure with adaptive variation, or to overcorrect for population structure and as a result, overlook adaptive markers, particularly for species whose demographic history is aligned with environmental gradients. In this case, the postglacial expansion of lodgepole pine likely matches the strong latitudinal gradient of winter temperatures. Since the analyses identifying GEA and GPA SNPs both adjusted for population structure, we may have eliminated some loci involved in local adaptation from consideration through this adjustment.

### 4.2 Climatic drivers of local adaptation

To design an assisted gene flow strategy that matches populations with suitable sites based on current and near-future climates, it is important to understand the climatic factors that have driven local adaptation. Once the key climatic factors for local adaptation are identified, a climate distance metric can be constructed to match seed sources with sites (e.g., Climate-Based Seed Transfer, O’Neill 2017, and Seedlot Selection Tool, https://seedlotselectiontool.org/sst/). Our GEA results for individual climate variables ranked the variable importance similarly to those identified based on growth in a 20-year field provenance trial and, to a lesser extent, to our seedling common garden phenotypes. Both sets of phenotypic data identified winter temperature variables including mean coldest month temperature, degree days below 0°, and extreme minimum temperature as important drivers of local adaptation. Other studies of these provenances (e.g., Liepe et al. 2016) and other populations of lodgepole pine in western Canada (e.g., Rweyongeza et al. 2007; Wang et al. 2010; McLane et al. 2011) corroborate these climatic variables as strong historic drivers of adaptation and differentiation, and at relatively broad spatial scales (Liepe et al. 2016). Nevertheless, the result that our set of neutral markers produced nearly equivalent climate variable rankings to the GEA set (Figure 4 vs. Figure S7) indicates that the substitution of genomic for phenotypic data needs to be approached with some caution.

Future pressures from drought are expected to become increasingly relevant for lodgepole populations as climate change progresses throughout the next century (Monserud et al. 2006, 2008; McLane et al. 2011). GEA-climate relationships were stronger than field phenotype-climate relationships for summer precipitation-related variables such as mean summer precipitation and cumulative moisture deficit (Figure 4). This suggest that water availability might result in diversifying selection across populations. A previous study with these populations found no significant population variation for drought-related seedling traits including stable carbon isotope ratios and biomass allocation to roots (Liepe et al. 2016); however, it did not include populations from drier provenances in the southern portion of the species range, and these may show stronger drought adaptation. Interestingly, the seedling common garden phenotypes in this study from the mild, maritime Vancouver climate had stronger associations with summer heat-related variables than the genomic data (Figure 4a).

None of the phenotypes we analyzed represent lifetime fitness. Nonetheless, the concordance of climatic drivers of seedling phenotypes, 20-year growth in the field, and genomic data are encouraging (Figure 4). The closest proxy to fitness among the seedling traits we analyzed may be seedling shoot mass, as a measure of growth during the common garden experiment. Trees that achieve larger sizes within the available frost-free period for growth will generally have higher fecundity as they have larger crowns with more sites for pollen and seed cone production. Forest managers are also ultimately interested in tree size for wood production, and trees with good juvenile growth are likely to grow well in a restoration context. We found weaker population differentiation for shoot mass than for the other seedling traits. Tree size is the product of many other component traits affecting seedling health and vigour, including phenology (which we analyzed directly as growth initiation and cessation), abiotic stress tolerance (including cold injury), resistance to insects and diseases, resource acquisition and allocation, physiological processes, cell density, etc. It is likely that loci underlying variation in growth have pleiotropic effects, and that they respond to selection through trade-offs in the various fitness consequences of component traits contributing to growth.

Which of these data sources – seedling phenotypes, field phenotypes, or genotypes – should be considered the standard against which the others are compared? One could argue that field-based growth over two decades better reflects meaningful provenance differences expressed in typical habitat. On the other hand, the precision phenotyping of seedlings for phenology and cold hardiness is difficult or impossible in long-term field trials, and these traits should be strongly linked with climate for boreal, sub-boreal, and montane species. Finally, it may be that the GEA-climate patterns provide the best indication of long-term selection as they may reflect periodic, episodic extreme climatic events causing injury and mortality that are not observed even over long field experiments. In any event, given the extensive overlap in top climate variables among these methods, we suggest that GEA approaches can rapidly provide information on climatic drivers of local adaptation for the design of assisted gene flow strategies when phenotypic data are not available. However, the potential for population structure to confound GEA approaches remains an important consideration.

### 4.3 Spatial scale of local adaptation to climate

We evaluated variation at adaptive loci against a model of localized versus rangewide genetic clines (Figure 1, *sensu* Barton 1999) along climatic temperature gradients (Figure 2). We found evidence of both localized and broad-scale genetic clines for clusters of SNPs associated with autumn cold injury (Figure 5 and Figure S8). Overall, the genetic clines associated with autumn cold injury do not exhibit the strongly sequential, localized clines envisioned by Barton (1999) and Savolainen et al. (2007), nor are all genetic clines coincident across the range of environments, but rather fit a model intermediate to the hypothetical scenarios illustrated in Figure 1B and 1C. Our study sampled provenances over only half of the species’ latitudinal range. It may be that sequential localized genetic clines would be more evident if our study included the full species range. While some clines for the major adaptive clusters we identified are largely variable across the range, there is a group of six SNPs that all show clines in the boreal region of the study area, but not in warmer areas (cluster 6 in Figure 5). These clines complement those of several other clusters for SNPs that are relatively invariant in the boreal portion of the range but vary in warmer regions (clusters 1, 2, and 4 in Figure 5). For instance, cluster 6 alleles conferring cold hardiness (the alternate PEA allele) have reduced standing variation in warmer provenances west of the Rocky Mountains, and follow both elevational and latitudinal patterns of temperature clines (Figures 5 and 6). Failure to detect polymorphisms for these SNPs in these provenances may be an artefact of small sample sizes (6<*n*<13) in a majority of the studied provenances (Figure S2). Nevertheless, these results indicate reduced genetic diversity for boreal-associated alleles of cluster 6. The absence of these alleles may be a limiting factor in seed transfer from sub-boreal to boreal climates, or across the Rocky Mountains. This localization may be indicative of alleles conferring additional cold hardiness in the coldest areas of the sampled range that may have trade-offs in the warmer areas (e.g., via pleiotropy or GxE such as conditional neutrality). Even so, the alleles in cluster 6 were not associated with the other phenotypes in our study (while all other clusters had associations to at least three phenotypes). Future investigation may be warranted, as the lack of pleiotropy inferred from associations to multiple phenotypes in cluster 6 may be a function of the cluster’s sample size, of linkage to unsampled antagonistic (regulatory) sites, conditional neutrality underlying gene action (or other GxE), of unmeasured phenotypes important to adaptation, or of other statistical and methodological shortcomings.

While our results suggest that localized genetic clines (Figure 5), and provenances associated with low genetic diversity in adaptive alleles (Figures 6 and S8), are evident in lodgepole pine, we did not find compelling evidence for localized genetic clines at scales that would constrain local seed transfer more narrowly than previous estimates of adaptive scales based on phenotypes (*cf.* Figure 4 in Liepe et al. 2016; Wang et al. 2010; Ukrainetz et al. 2018) or current seed transfer policy would suggest (Ying and Yanchuk 2006; O’Neill et al. 2017), nor at scales that would necessitate highly localized spatial genetic conservation units. At present, British Columbia’s genetic conservation program for forest trees uses British Columbia’s 16 Biogeoclimatic Ecological Classification (BEC) zones to assess adequacy of both *in situ* (Hamann et al. 2004; Chourmouzis et al. 2009) and *ex situ* (Krakowski et al. 2009) genetic conservation for all 50 of BC’s native tree species. If other species show patterns of distribution of adaptive diversity similar to lodgepole pine, continued management of conservation populations within these ecological zones should be sufficient (Liepe et al. 2016).

Climate-based local adaptation has long been observed for many plant species (Leimu and Fisher 2008; Hereford 2009) including conifers (Langlet 1971; Savolainen et al. 2007; Boshier et al. 2015; Lind et al. 2018), the scale of which is determined by the interplay between migration, selection, and drift (discussed in Lenormand 2002, Tigano & Friesen 2016). Historically, the spatial scales over which local adaptation occurs has been inferred from both short- and long-term transplant experiments (Langlet 1971; Morgenstern 1996). Only recently has the technology been available to study the spatial distribution of adaptive variation at loci across the genome. This new source of insight into local adaptation comes at a time when climate change creates an imperative for mitigating inevitable risks of productivity loss and threats to natural populations across forestry, agricultural, and natural systems. The common sources of data used towards such purposes, such as field provenance trials, seedling common gardens, scale-free spatial climatic data, and genomic studies, however, come with varied logistical limitations and are not always feasible or appropriate in every situation (Blanquardt et al. 2013; Sork et al. 2013; Gibson et al. 2016; Hoban et al. 2016; Flanagan et al. 2018). The large number of phenotyped and genotyped provenances in this study allow us to quantify and compare detailed spatial and climatic patterns of adaptive variation, and to assess their utility for planning assisted gene flow, the need for *in situ* and *ex situ* genetic conservation, and the potential for populations to adapt to new climates without intervention. While our data are for lodgepole pine, we hope these results will inform and accelerate climate adaptation efforts with other widespread species.

## Acknowledgements

This research used data generated by the AdapTree Project (S.N.A. and Andreas Hamann, coproject leaders), and was analyzed as part of the CoAdapTree Project. These projects were funded by Genome Canada, Genome BC, with co-funding from Alberta Innovates Bio Solutions, the BC Ministry of Forests, Lands and Natural Resources Operations, and the Forest Genetics Council of BC (a full list of sponsors is available at http://coadaptree.forestry.ubc.ca/sponsors/). Support for C.M. was also provided by a grant to S.N.A. from the Forest Genetics Council of British Columbia and an NSERC Discovery Grant to S.N.A. Seedlots were provided by 63 government agencies and forest companies (listed at http://adaptree.forestry.ubc.ca/seed-contributors/). Thanks to Kristin Nurkowski, Robin Mellway and Pia Smets for laboratory work including DNA extraction and phenotyping. The lodgepole pine SNP array was designed based on analyses by Sam Yeaman, Kay Hodgins, and Katie Lotterhos, with assistance and support from Affymetrix. The BC Ministry of Forests, Lands and Natural Resource Operations provided data from the Illingworth Provenance Trial. Additional technical assistance and suggestions for this manuscript were provided by members of the AdapTree and CoAdapTree teams and Aitken lab members. We thank Sam Yeaman for providing comments on an earlier version of this manuscript.

## Author contributions

SNA and CRM conceived this project. Common garden experiments and phenotyping by IRM. SNP tables compiled by JBY and IRM. Analyses designed by CRM with input from IRM, SNA, and BML. GPA by IRM. GEA by BML. Figures by CRM. Analysis of Illingworth trial by TW. Introduction by SNA, BML, CRM and JBY. Methods by IRM and CRM. Results by CRM. Discussion by CRM, SNA, BML and JBY. Supporting information document by CRM.

## Archiving statement

Data for this study will be made publicly available upon acceptance.

## Supplementary Information

**Table S1.** Overlap between genotype-environment associations (GEA) from bayenv2 and genotype-phenotype associations (GPA). The GEAs column indicates the number of environmentally associated loci that met our filtering criteria (locus in top 300 ranks for Bayes factor across all three chains, and also in top 300 ranks for *rho* across all three chains, after filtering for one SNP per contig). Numbers in phenotypic columns indicate the overlap of GEAs with GPA hits (i.e., the same position in each association), with the number in the parenthesis being the additional number of unique GPA hits that were within 1000bp of a GEA hit. Unique GPA Hits is the unique number of loci across these phenotypic columns for a given environmental variable, with parenthetical values indicating the additional number of unique GPA hits within 1000bp of a GEA. The last row indicates the unique number of SNPs from a given column. Environmental abbreviations as in Table 1 of main text.

| Environmental Variable | GEAs | Budbreak | Budset | Cold injury | Shootmass | Unique GPA Hits |
| --- | --- | --- | --- | --- | --- | --- |
| AHM | 202 | 15 (5) | 41 (23) | 4 (2) | 1 (1) | 52 (26) |
| CMD | 238 | 32 (12) | 67 (33) | 36 (12) | 25 (8) | 86 (36) |
| DD5 | 192 | 46 (26) | 32 (11) | 48 (14) | 39 (9) | 82 (32) |
| DD_o | 142 | 36 (21) | 35 (21) | 37 (15) | 32 (10) | 63 (32) |
| EMT | 129 | 49 (17) | 30 (12) | 42 (12) | 32 (9) | 69 (26) |
| EXT | 26 | 12 (4) | 14 (6) | 15 (4) | 15 (3) | 15 (6) |
| Eref | 234 | 37 (9) | 69 (29) | 42 (14) | 30 (9) | 93 (35) |
| FFP | 193 | 65 (17) | 31 (7) | 50 (9) | 39 (8) | 95 (23) |
| MAP | 117 | 19 (11) | 31 (23) | 7 (5) | 2 (3) | 43 (29) |
| MAT | 186 | 59 (30) | 40 (25) | 44 (16) | 35 (10) | 93 (47) |
| MCMT | 188 | 49 (22) | 29 (14) | 41 (14) | 30 (9) | 70 (30) |
| MSP | 202 | 23 (10) | 52 (32) | 16 (9) | 10 (9) | 64 (35) |
| MWMT | 141 | 21 (8) | 35 (12) | 41 (12) | 39 (8) | 56 (18) |
| NFFD | 2 | 0 (0) | 0 (0) | 0 (0) | 0 (0) | 0 (0) |
| PAS | 112 | 0 (1) | 3 (3) | 4 (1) | 3 (1) | 9 (4) |
| SHM | 233 | 30 (10) | 65 (32) | 33 (11) | 22 (9) | 85 (35) |
| TD | 161 | 10 (9) | 9 (13) | 15 (10) | 7 (7) | 19 (17) |
| bFFP | 167 | 52 (22) | 26 (9) | 43 (13) | 37 (10) | 79 (30) |
| eFFP | 174 | 64 (18) | 29 (8) | 45 (11) | 35 (10) | 90 (26) |
| unique loci | 901 | 105 (44) | 102 (63) | 77 (34) | 49 (24) | 212 (105) |

**Figure S1:**
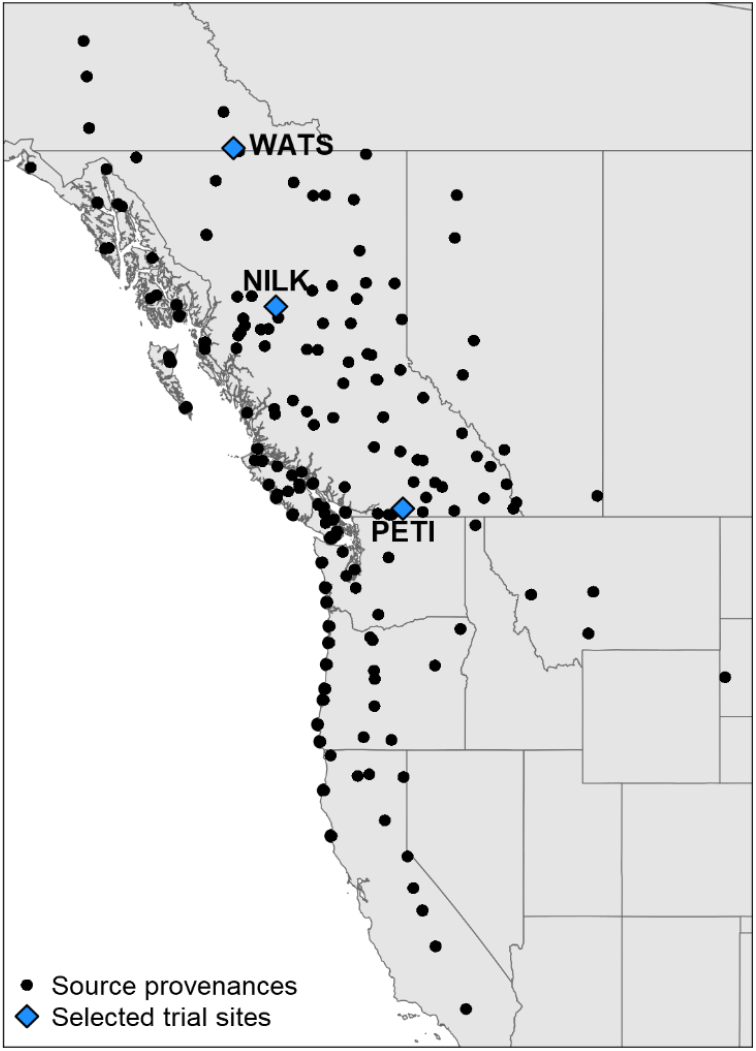
Source provenances and selected trial sites for the Illingworth lodgepole pine provenance trial.

**Figure S2:**
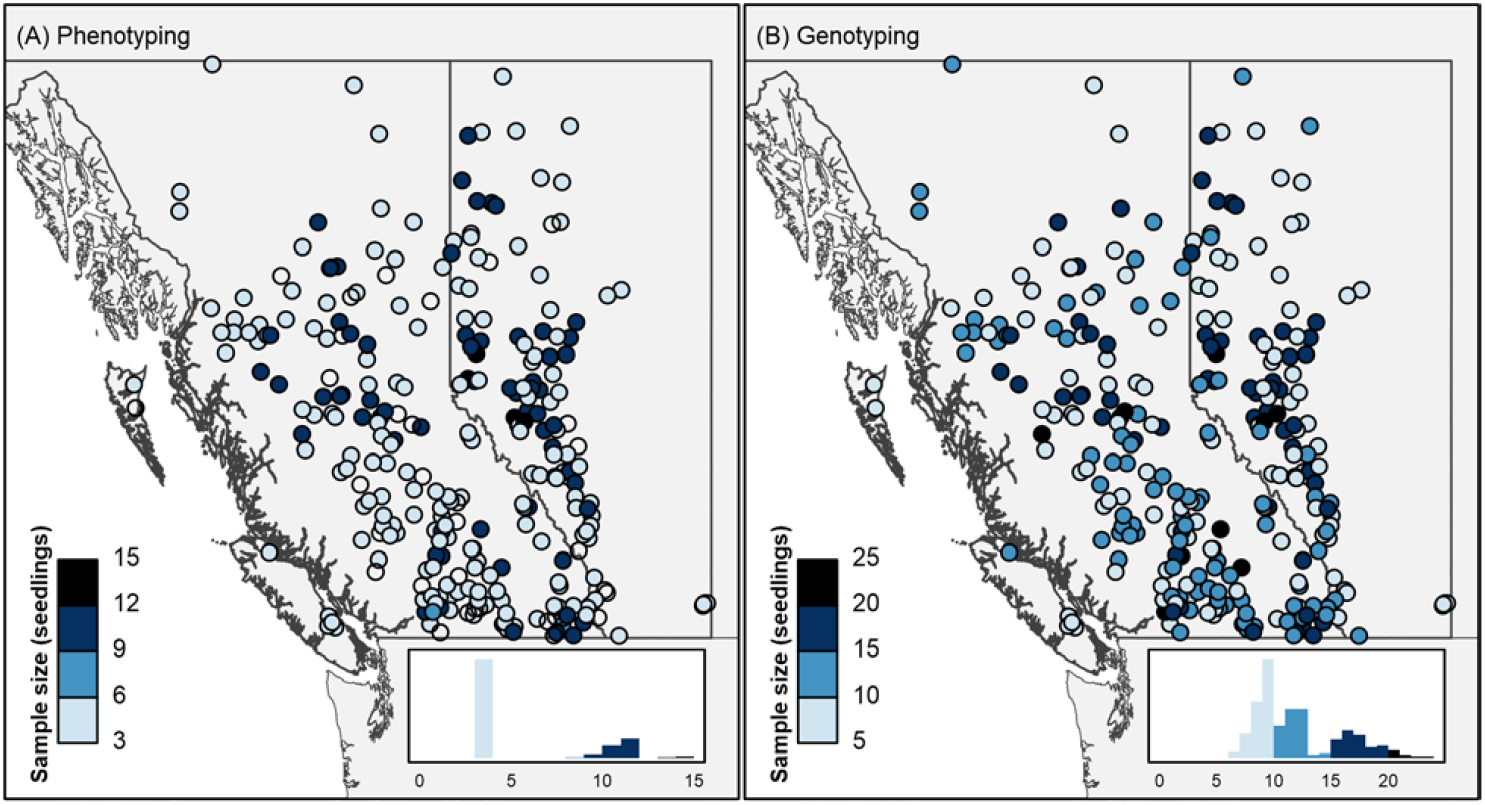
Sample size of common garden phenotyping and genotyping. SNP array Genotyping was conducted on phenotyped common garden seedlings and an additional sample of seedlings grown in a growth chamber.

**Figure S3:**
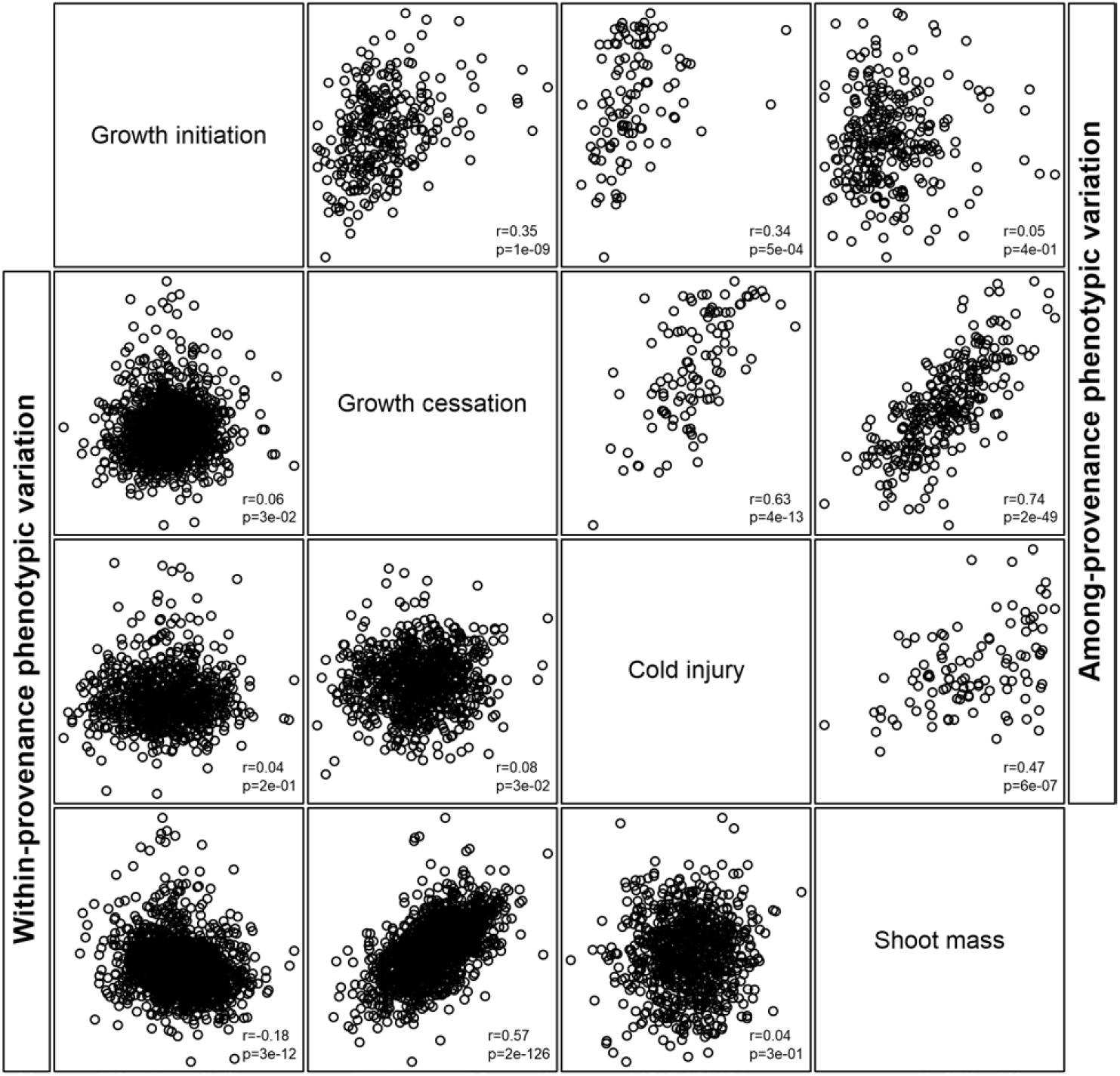
Among- and within-provenance relationships between the four traits. Among-provenance variation is the variation of provenance-mean phenotypic values. Within-provenance variation is the variation of individual seedling phenotypes that have had their provenance-mean phenotypic value subtracted.

**Figure S4:**
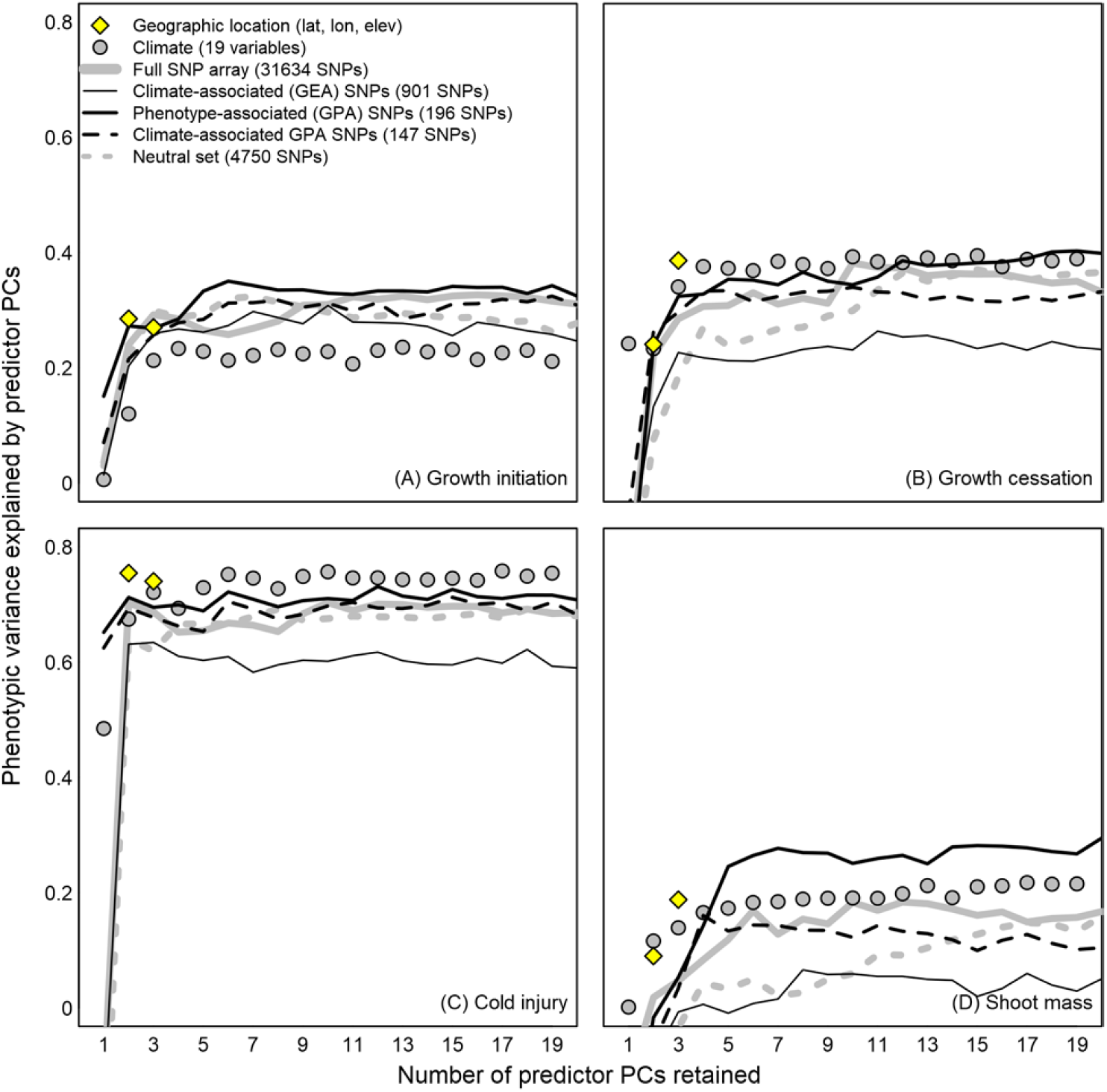
Equivalent analysis for Figure 3, using Random Forest regression instead of linear regression. Each point is the pseudo-*R*^2^ of a Random Forest regression of provenance-mean phenotype against the specified number of principal components of the predictor data. GEA SNPs (thin black line) are the pooled top-300 SNPs based on Bayes factor from each of the 19 climate variables. GPA SNPs (thick black line) are the top 1% of coding-region SNPs (maximum of one SNP per contig) based on the p-value of a population-structure-corrected linear association of allele frequencies to seedling phenotypes. Climate-associated GPA SNPs (black dashed line) are GPA SNPs with further support for strong association to climate (see methods). The neutral set is shown as a grey dashed line

**Figure S5:**
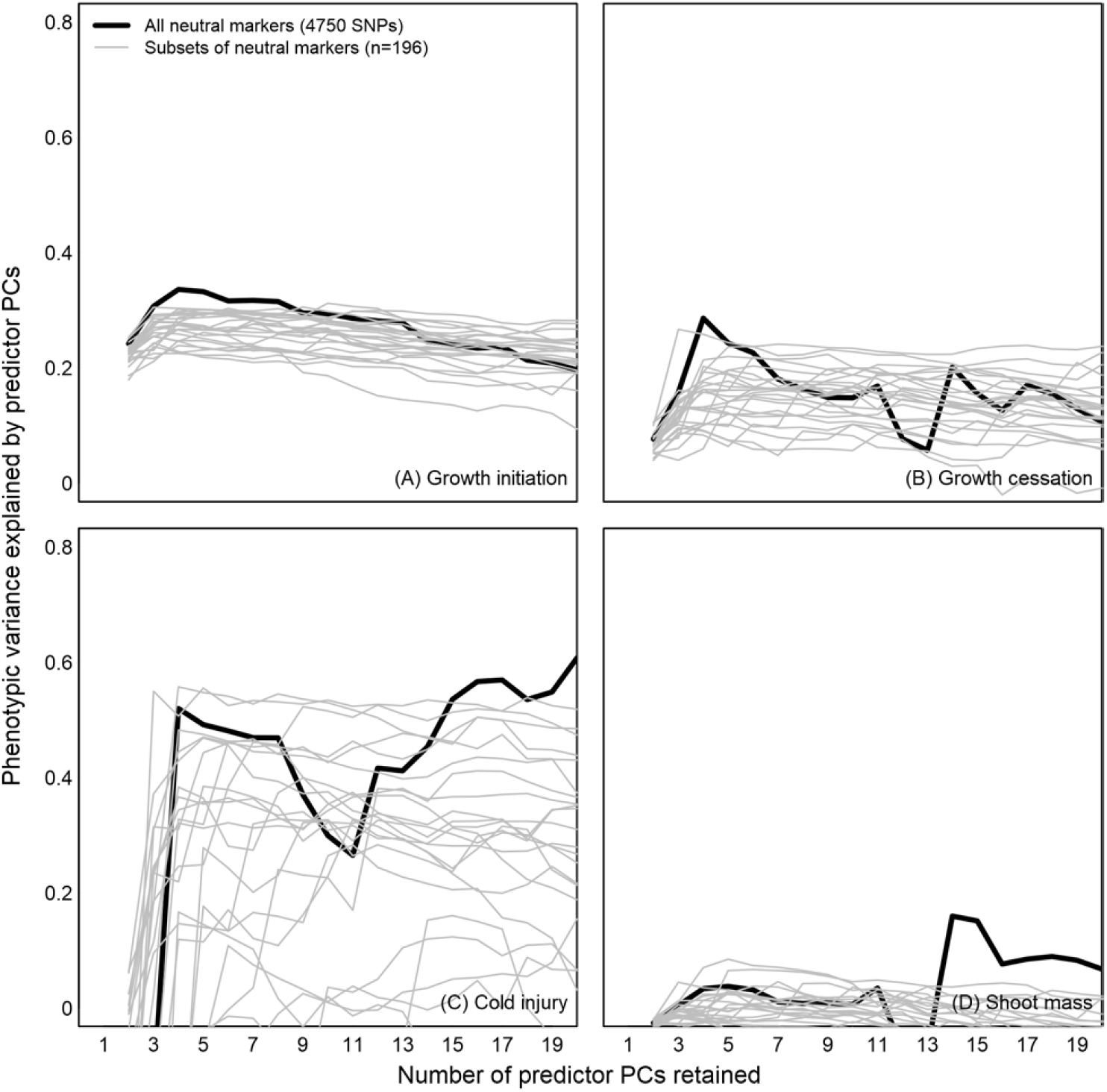
Explanatory power of small subsets of neutral SNPs. Each point is the cross-validated ***R**^2^* of a multiple linear regression of provenance-mean phenotype against the specified number of principal components of minor allele frequency in an n=196 subset of neutral SNPs. Each grey line is a different subset, selected sequentially from the neutral set. The black line is the equivalent analysis for the full neutral set.

**Figure S6:**
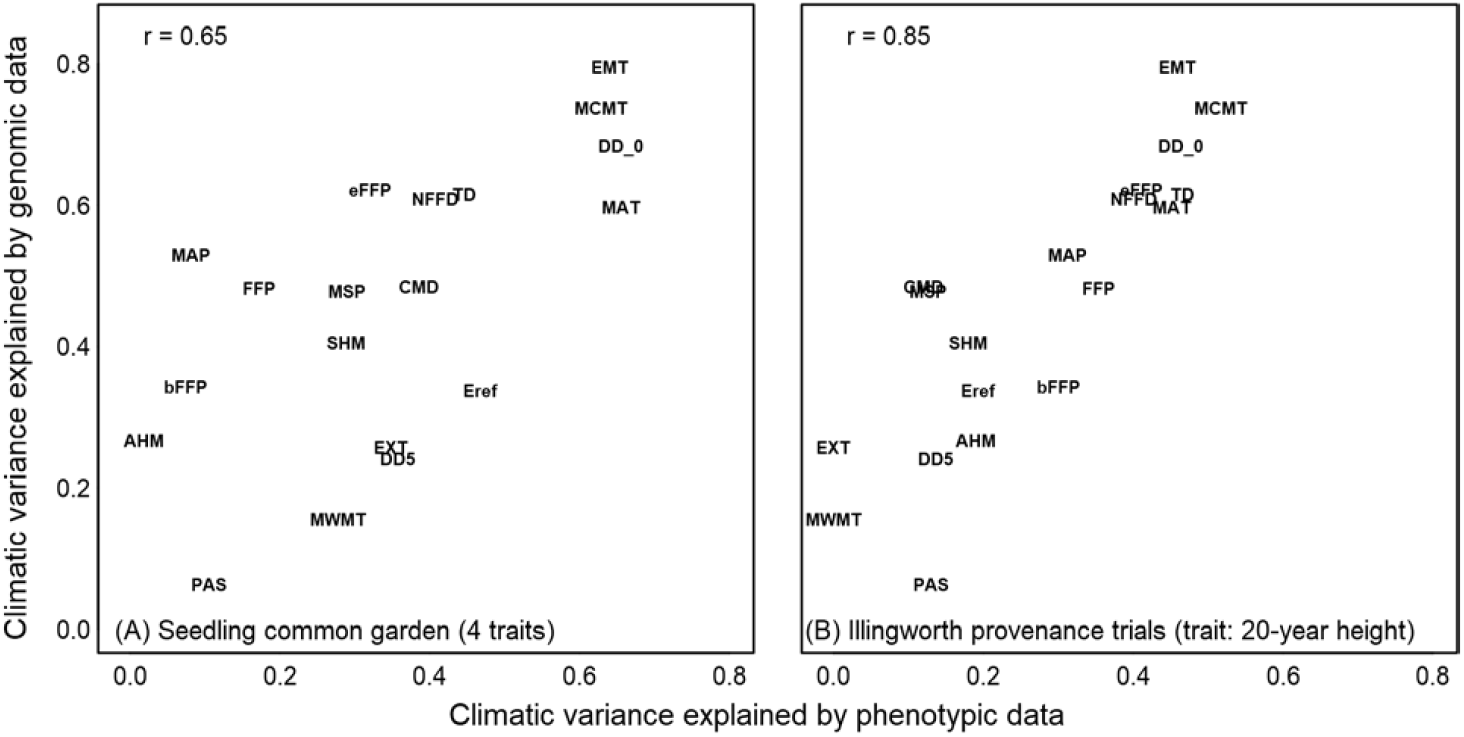
Climatic variable selection based on phenotypic vs genomic data, equivalent to Figure 4 except using the full SNP array instead of GEA SNPs. Variance explained is the cross-validated *R*^2^ of a multiple linear regression of each climate variable (response variable) against the phenotypic or genomic predictor variable set. Genomic data (predictor variables for the y-axis analyses) are four principal components of the minor allele frequencies for the full SNP array (n=31634 SNPs). Phenotypic data (predictor variables for the x-axis analyses) for panel A are provenance-mean phenotypes for the four common-garden traits presented in Figure 2. Phenotypic predictor data for panel B are 20-year heights of the Illingworth lodgepole pine provenance trial. Climate variable acronyms are described in Table 1.

**Figure S7:**
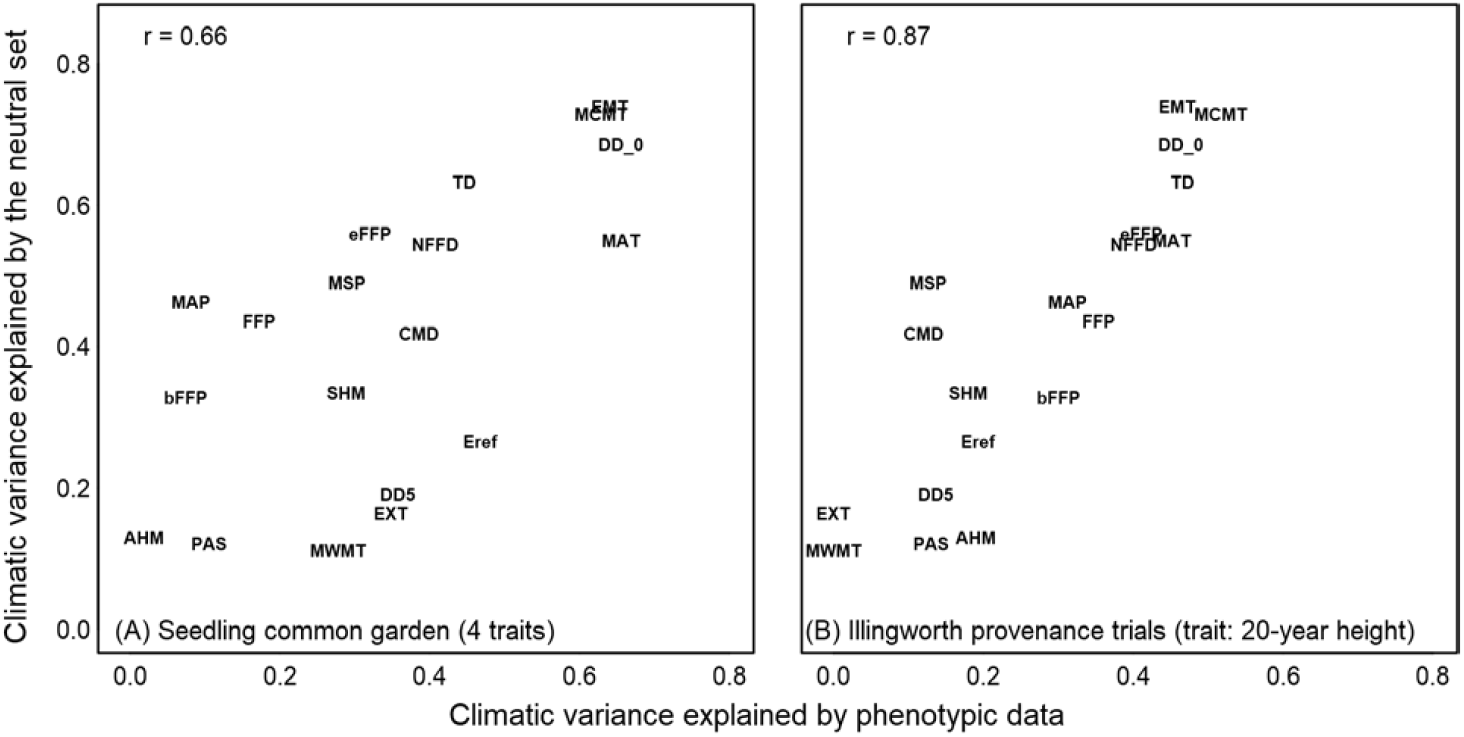
As in Figure S6 above, but using the neutral set (n=4750 SNPs) instead of the full SNP array.

**Figure S8:**
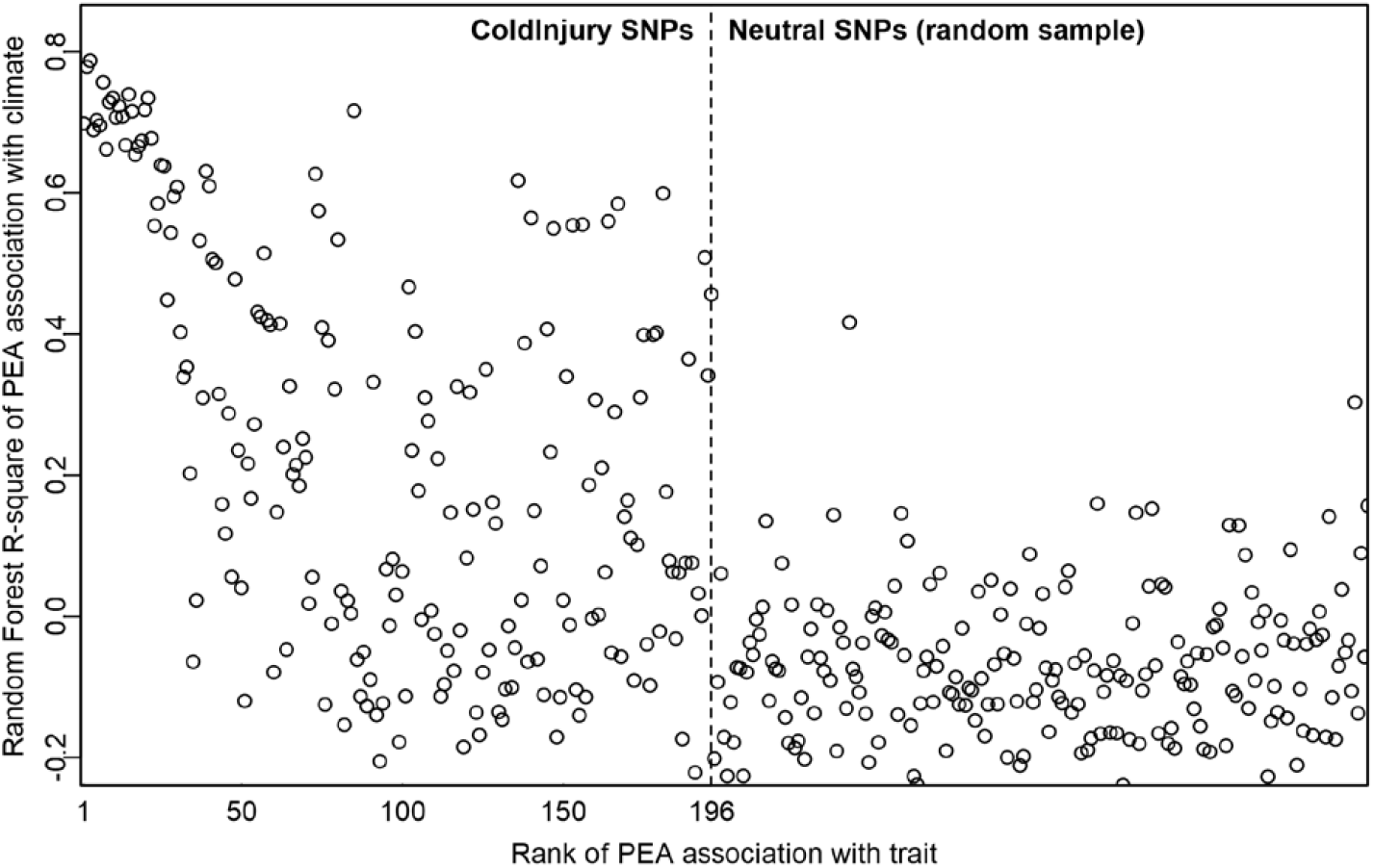
Relationship of GPA-selected loci to climate. The y axis is the pseudo-*R*^2^ of a random forest regression of population-mean PEA frequency (response variable) to the 19 bioclimate predictor variables. PEAs are arranged in order of increasing GPA p-value (decreasing significance), with a random sample of neutral SNPs shown for comparison.

**Figure S9:**
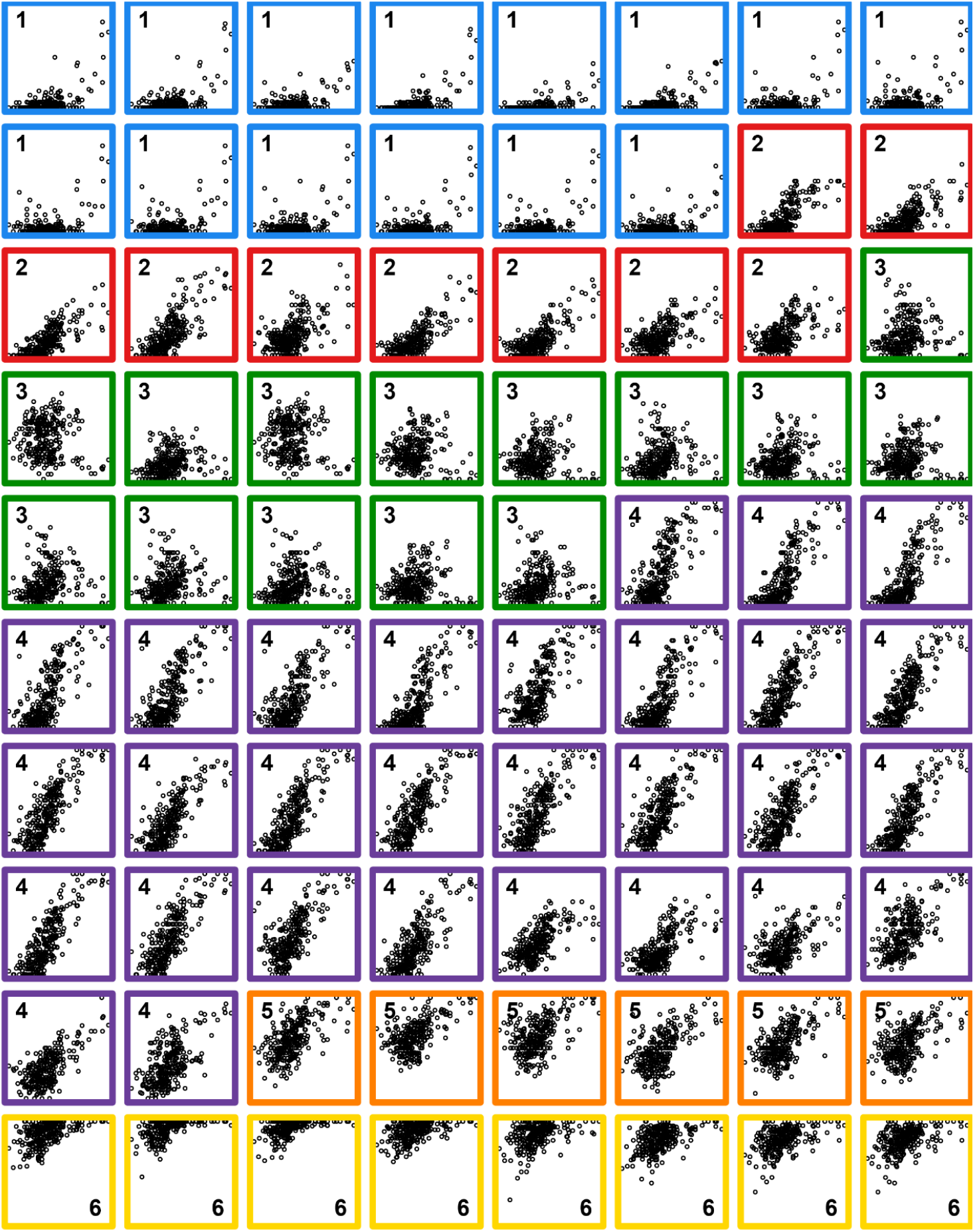
Genetic clines of climate-associated (pseudo-ß^2^ > 0.32) GPA loci for autumn cold injury. Loci are clustered by PEA frequency across provenances. The x axis is autumn mean daily minimum temperature; the y axis is population-mean PEA frequency.

**Figure S10:**
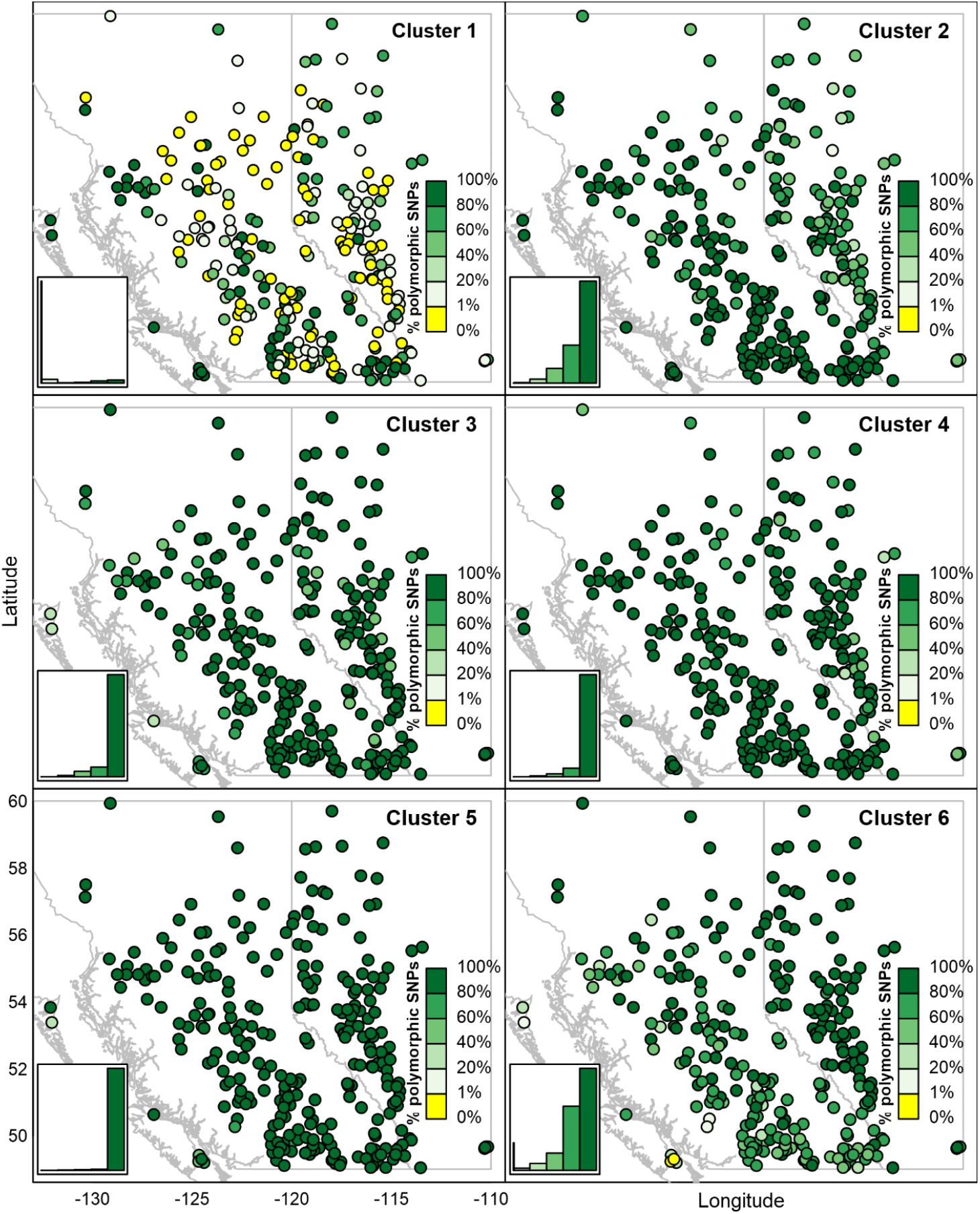
proportional polymorphism, by cluster, for each provenance: The percentage of the SNPs in each cluster that have standing variation in both alleles, i.e., ***H_e_*** > 0.

**Figure S11:**
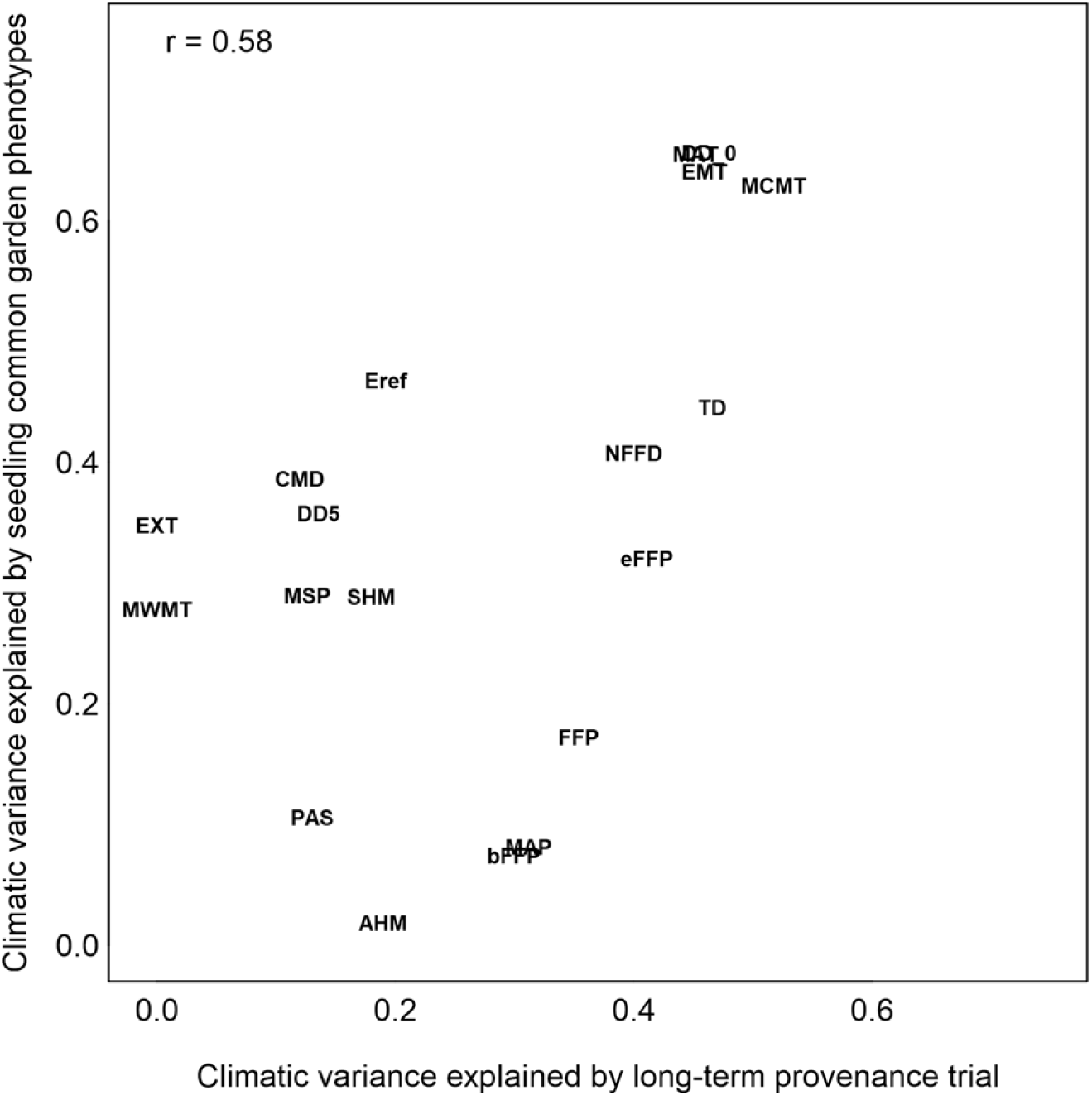
Climatic variable selection based on seedling common garden phenotypes vs. long-term provenance trial heights. Variance explained is the cross-validated *R*^2^ of a multiple linear regression of each climate variable (response variable) against the phenotypic predictor variable set. Phenotypic predictor data for the x-axis are 20-year heights of the Illingworth lodgepole pine provenance trial. Predictor variables for the y-axis are provenance-mean phenotypes for the four common-garden traits presented in Figure 2. Climate variable acronyms are described in Table 1.

**Figure S12:**
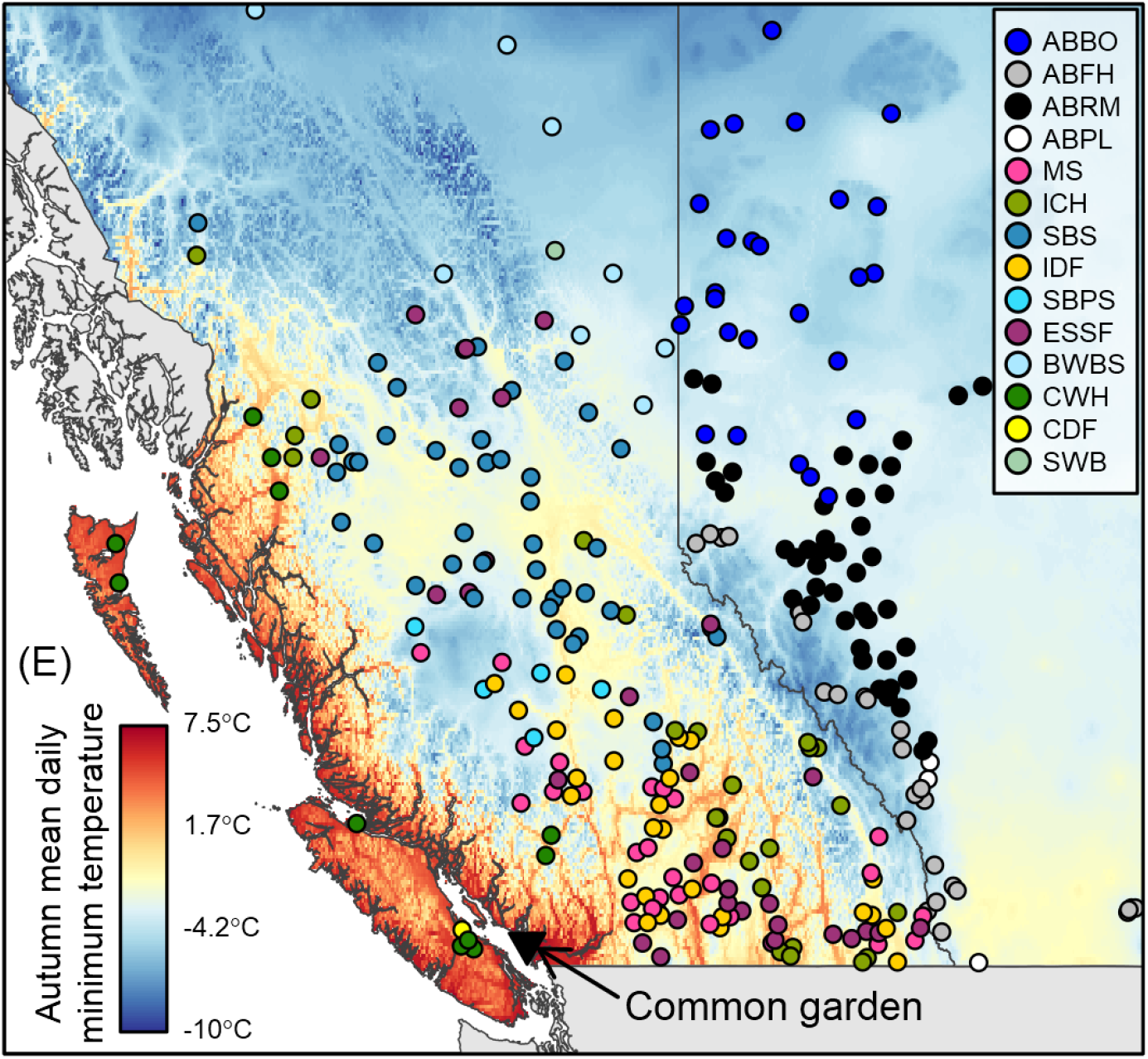
Biogeoclimatic zones (British Columbia) and natural regions (Alberta) of each sampled provenance.

**Figure S13:**
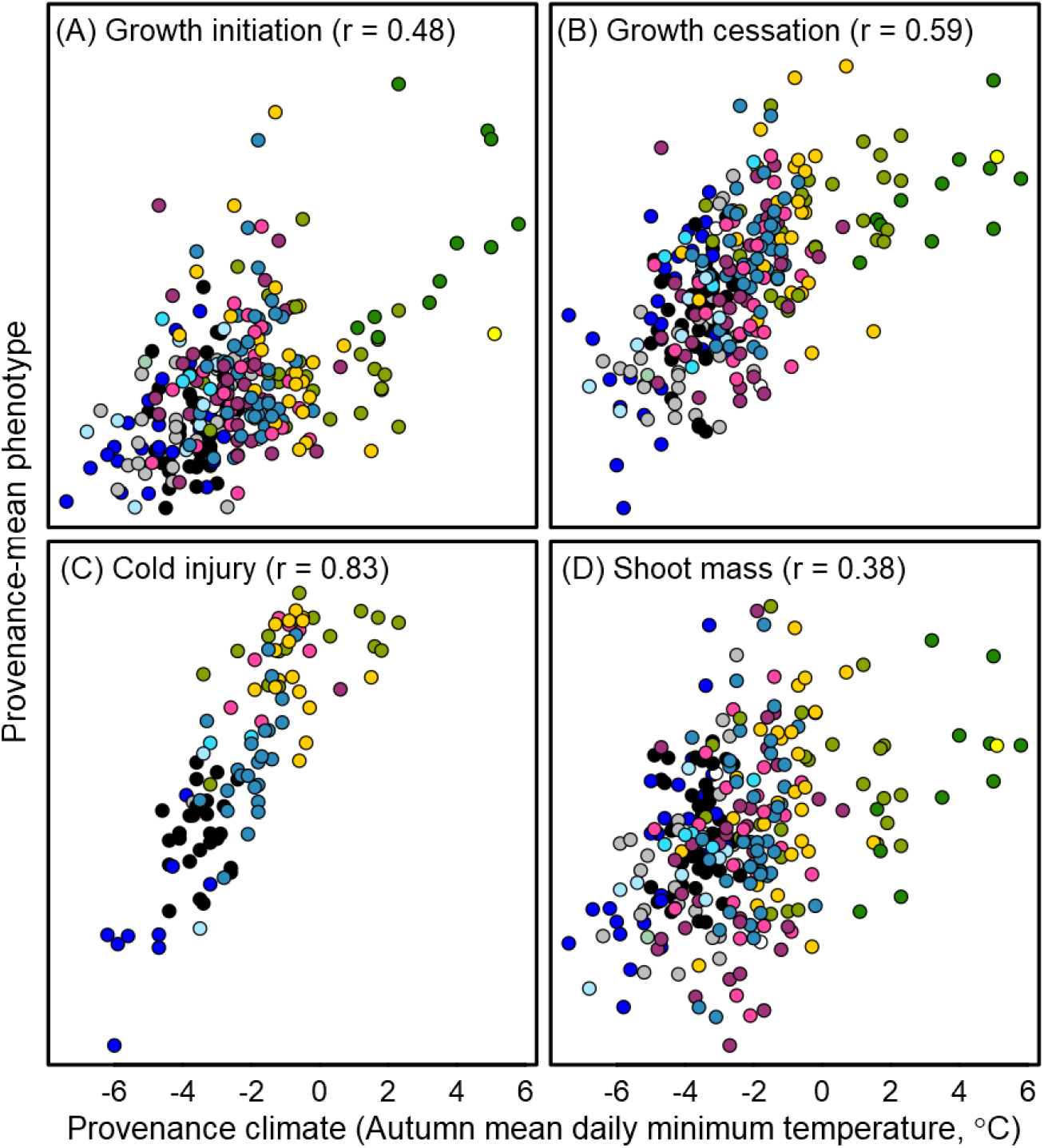
**Phenotypic clines of four traits in lodgepole pine seedlings grown in the Vancouver common garden**, colour themed by biogeoclimatic zone (British Columbia) and natural region (Alberta). See Figure S12 for map and color schemes.

**Figure S14:**
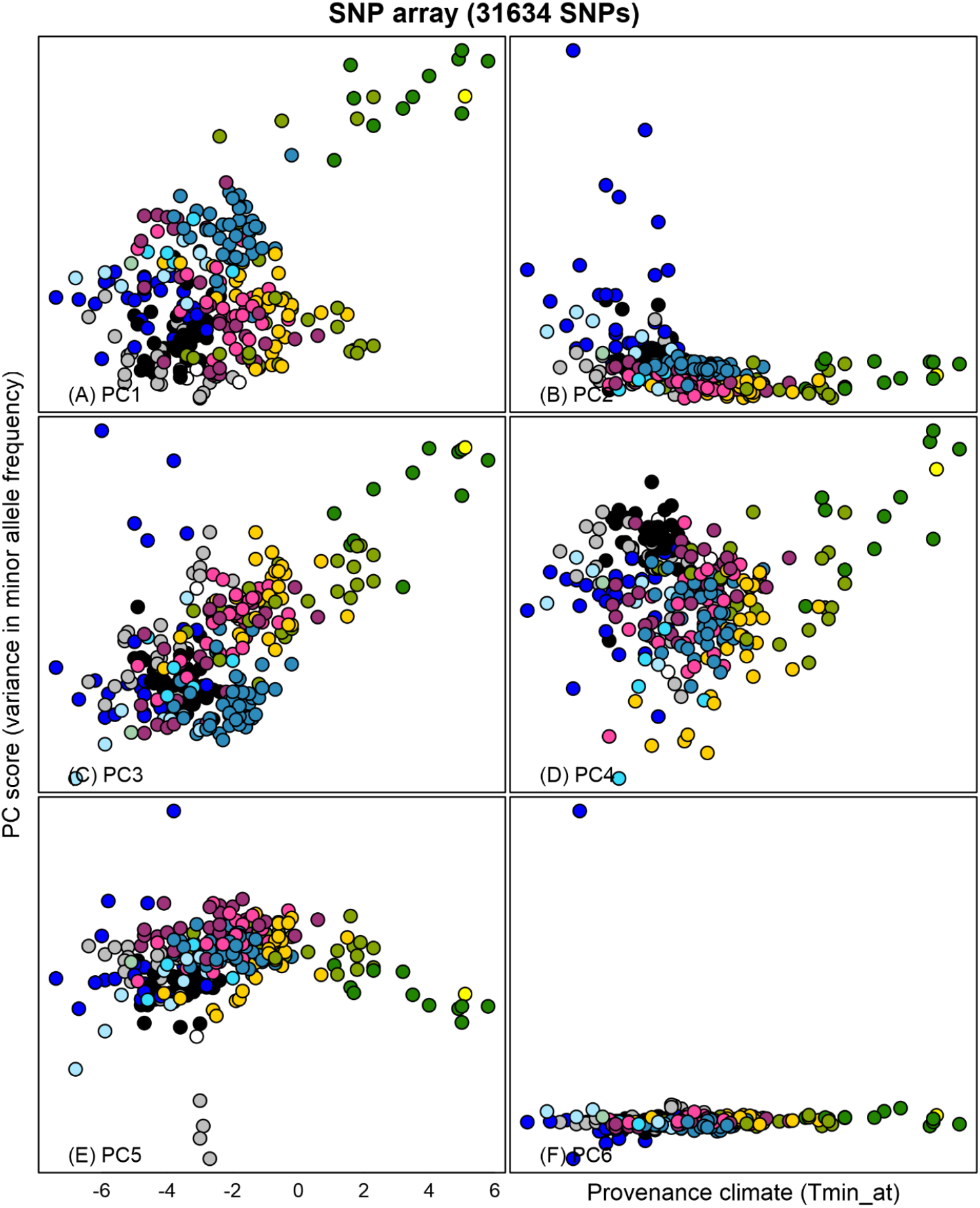
**First six principal components of *z*-standardized provenance-mean minor allele frequencies in the full SNP array** (excluding the neutral set), plotted against autumn temperature. Provenances are colour themed by biogeoclimatic zone (British Columbia) and natural region (Alberta). See Figure S12 for map and color scheme.

**Figure S15.**
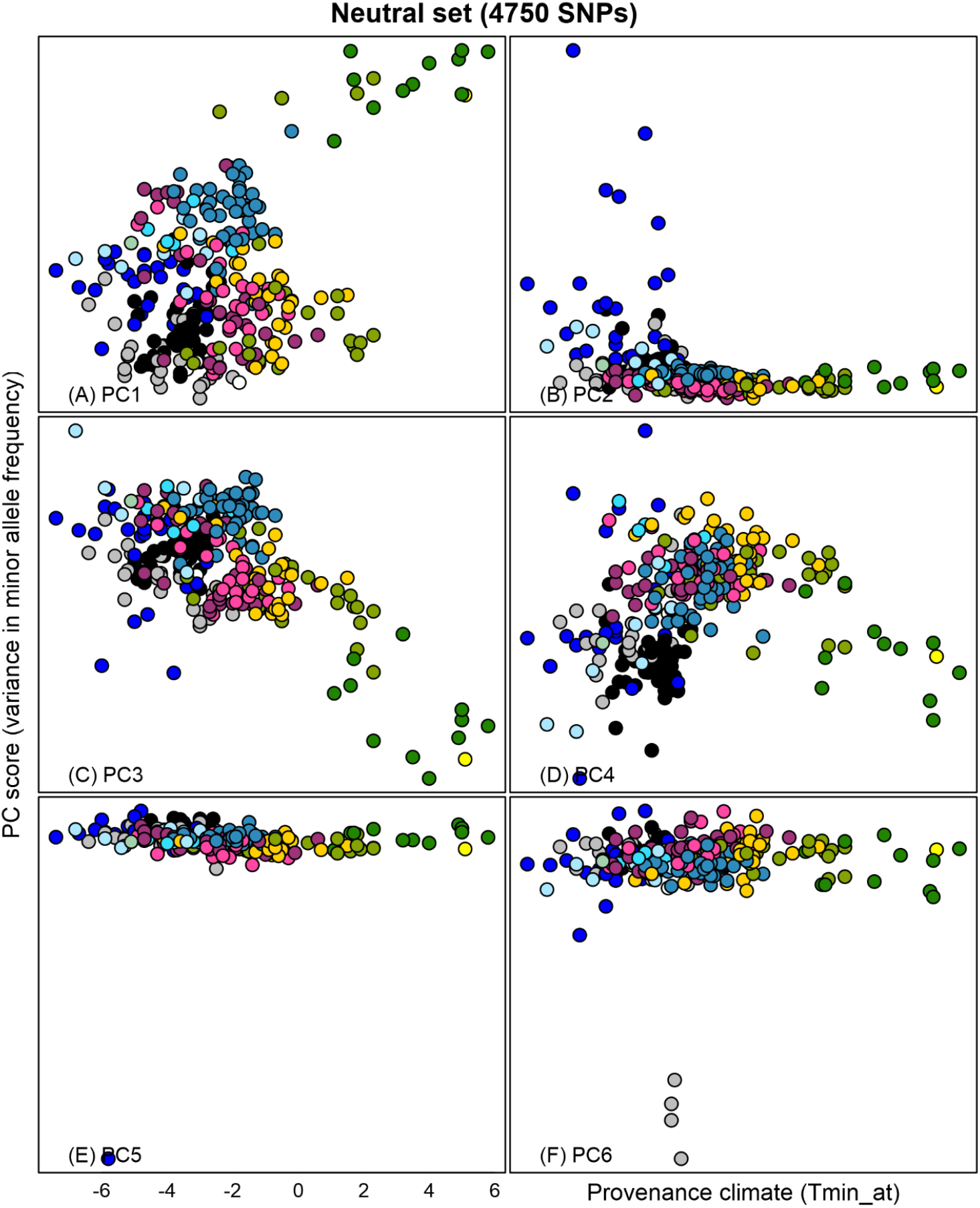
**First six principal components of *z*-standardized provenance-mean minor allele frequencies in the neutral set**, plotted against autumn temperature. Provenances are colour themed by biogeoclimatic zone (British Columbia) and natural region (Alberta). See Figure S12 for map and color scheme.

**Figure S16.**
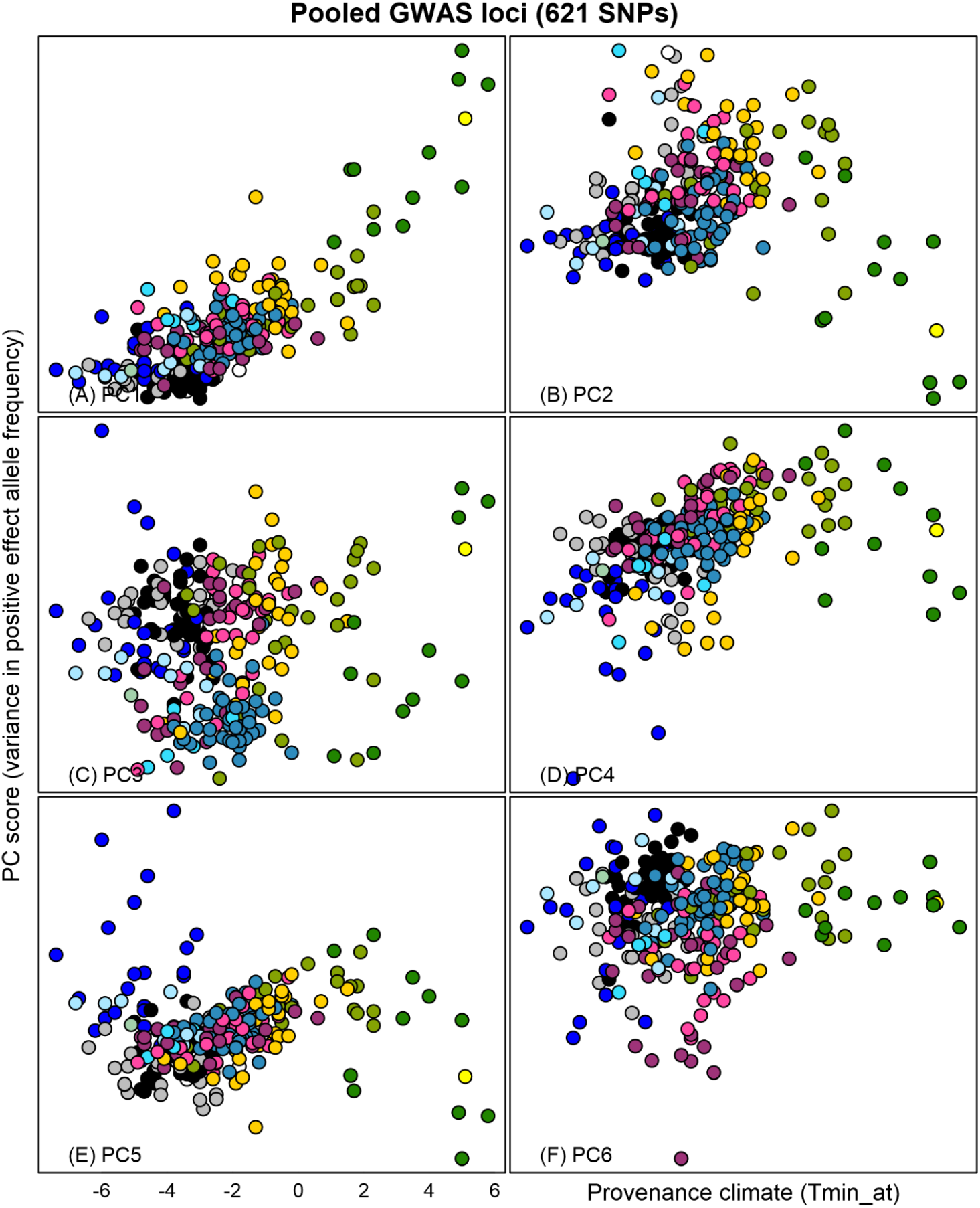
**First six principal components of *z*-standardized provenance-mean positive-effect allele frequencies in the pooled GWAS loci for all four common garden traits**, plotted against autumn temperature. Provenances are colour themed by biogeoclimatic zone (British Columbia) and natural region (Alberta). See Figure S12 for map and color scheme.

